# High-throughput mRNA-seq atlas of human placenta shows vast transcriptome remodeling from first to third trimester

**DOI:** 10.1101/2023.06.06.543972

**Authors:** Tania L Gonzalez, Sahar Wertheimer, Amy E Flowers, Yizhou Wang, Chintda Santiskulvong, Ekaterina L Clark, Caroline A Jefferies, Kate Lawrenson, Jessica L Chan, Nikhil V Joshi, Yazhen Zhu, Hsian-Rong Tseng, S. Ananth Karumanchi, John Williams, Margareta D Pisarska

## Abstract

**Background:** The placenta, composed of chorionic villi, changes dramatically across gestation. Understanding differences in ongoing pregnancies are essential to identify the role of chorionic villi at specific times in gestation and develop biomarkers and prognostic indicators of maternal-fetal health.

**Methods:** The normative mRNA profile is established using next-generation sequencing of 124 first trimester and 43 third trimester human placentas from ongoing healthy pregnancies. Stably expressed genes not different between trimesters and with low variability are identified. Differential expression analysis of first versus third trimester adjusted for fetal sex is performed, followed by a subanalysis with 23 matched pregnancies to control for subject variability using the same genetic and environmental background.

**Results:** Placenta expresses 14,979 mRNAs above sequencing noise (TPM>0.66), with 1,545 stably expressed genes across gestation. Differentially expressed genes account for 86.7% of genes in the full cohort (FDR<0.05). Fold changes highly correlate between the full cohort and subanalysis (Pearson = 0.98). At stricter thresholds (FDR<0.001, fold change>1.5), there are 6,941 differentially expressed protein coding genes (3,206 upregulated in first and 3,735 upregulated in third trimester).

**Conclusion:** This is the largest mRNA atlas of healthy human placenta across gestation, controlling for genetic and environmental factors, demonstrating substantial changes from first to third trimester in chorionic villi. Specific differences and stably expressed genes may be used to understand the specific role of the chorionic villi throughout gestation and develop first trimester biomarkers of placental health that transpire across gestation, which can be used for future development of biomarkers in maternal-fetal disease.

## Introduction

The placenta consists of chorionic villi that are derived from the trophoblast cells of the embryo. Following implantation and maternal signaling, placentation is established in the late first trimester (1). Throughout gestation, placenta serves a dynamic and critical role that impacts both maternal and fetal outcomes and provides communication between the fetal and maternal systems while providing the fetus with nutrients and removing waste (2). Additionally, the placenta has metabolic and endocrinologic functions which serve to regulate activity in pregnancy as well as to protect the fetus from exposures (1–3). However, molecular features of the placenta in the late first trimester differ considerably from the third trimester placenta, despite a similar tissue architecture (2). Gene expression changes likely correlate with functional changes at different gestational ages and suggest the placenta undergoes complex rearrangements to support fetal development, as suggested by microarray studies (4–8). Understanding the normative placental transcriptome signature, including stably and differentially expressed genes throughout gestation, may aid in understanding healthy placental function and serve as a comparison for alterations that drive pathology (9–12). Next generation sequencing technologies, such as RNA sequencing (RNA-seq), allows for a more comprehensive view of the entire placental transcriptome including the identification of novel transcripts compared to older methods such as microarrays (13). Although there are a few RNA-seq studies comparing the first trimester placenta to later gestational ages, the studies are small with only 5-8 subjects at first trimester, and use chorionic villi collected from pregnancy terminations which may have abnormal placentation, not reflective of a healthy placenta in ongoing pregnancies (14, 15). Furthermore, subject variability and underlying maternal and fetal conditions may contribute to differences that are not indicative of the normative state. To define the normative placenta in the first trimester, a large cohort of placenta from ongoing healthy pregnancies is needed. Furthermore, to control for environmental and genetic factors, and minimize subject variability, corresponding third trimester placenta for the same subjects are needed to define the normative human placenta transcriptome as pregnancy progresses.

We developed the largest mRNA-seq study that includes the most first trimester placentas to date (n=124), including 23 subjects sequenced at both first and third trimester, which controls for genetic, demographics, and environmental variability among different subjects. Additionally, first trimester tissue was obtained from chorionic villus sampling, not terminations, allowing us to confirm normal placenta development leading to a healthy pregnancy and delivery. Our normative human placental transcriptome as gestation progresses, with differentially and stably expressed genes may be used to develop first trimester biomarkers of placental health that transpires across gestation, which will be the foundation for future development of biomarkers in maternal-fetal disease.

## Materials and Methods

### Study Population

The study population consisted of 144 singleton spontaneous pregnancies between 2009 and 2018, including 124 with first trimester (59 female, 65 male) and 43 with third trimester (18 female, 25 male) placentas available. Subjects with infertility, pre-existing diabetes or hypertension were excluded. All protocols were performed with informed consent in accordance with the institutional review board’s guidelines at the Cedars-Sinai Medical Center under IRB protocols Pro00006806 and Pro00008600. All pregnancies resulted in the delivery of a viable infant.

### Analysis of demographic data

Demographics of pregnancies from the first versus third trimester placenta sequencing groups were compared. Specified pregnancy complications included pregnancy-induced hypertension, gestational diabetes, coagulation disorders, placenta previa, and placental abruption. Three patients classified as “placenta other” had either low-lying placenta, adherent placenta or velamentous cord insertion. Means and standard deviations were reported for continuous variables. Proportions were reported as percentages. Student’s T-test was used for normally distributed continuous variables, and the Wilcoxon rank-sum test was used for non-parametric data. Fisher’s exact test was used when appropriate. For comparison of categorical variables, the Chi-square test was used.

### Collection of placenta samples

Placental research samples were collected from tissue which is normally discarded after clinical visits. First trimester placenta samples were collected between 70- and 100-days gestation after CVS procedures for prenatal diagnosis. Fetal-derived chorionic villi (first trimester placenta tissue) were cleaned and separated from any maternally-derived decidua (non-placenta tissue), and 5-25 mg were used for research. Third trimester placenta samples were collected between 254- and 290-days gestation, after delivery of a viable neonate. One centimeter cubed of placental tissue samples were obtained immediately after delivery from the fetal side of the placenta near the site of cord insertion beneath the amnion. Tissue samples were kept on ice and submerged in RNA*later* RNA stabilization reagent (QIAGEN, Hilden, Germany) within 30 minutes of collection and stored at -80°C.

### RNA extraction from first trimester placenta

RNA extraction was performed from the first trimester placenta samples utilizing a method optimized for delicate tissue (16–18). Briefly, tissue samples were thawed on ice with 600 µl of RLT Plus lysis buffer (QIAGEN) and 1% β-mercaptoethanol added to each sample. Tissue was homogenized by passing at least 10 times through needles (22G, 25G, and 27G) attached to an RNase-free syringe. Homogenates were loaded onto AllPrep spin columns and further processing followed manufacturer instructions for the AllPrep DNA/RNA/miRNA Universal Kit (QIAGEN). RNA was eluted with 30-50 µl of RNase-free water at room temperature and the elution was passed through the column twice to improve yields, as previously described (16–18). For the 124 samples analyzed, the average RNA integrity number was 8.81.

### RNA extraction from third trimester placenta

Third trimester placental tissue was thawed on ice, then a quarter of collected tissue was diced with single use blades coated in RNA*later* buffer. Tissue was sonicated on ice in lysis buffer (600 µl RLT Plus lysis buffer with 1% beta-mercaptoethanol buffer) using 5 second pulses on a low setting (#2 on the Branson Sonifier 150D, CT, USA) until tissue fragments were small enough to pass through needles and continue RNA isolation with the first trimester method. For the 43 samples analyzed, the average RNA integrity number was 8.82.

### Library preparation and mRNA sequencing

Library construction was performed using the Illumina TruSeq Stranded mRNA library preparation kit (Illumina, San Diego, CA). Briefly, total RNA samples were assessed for concentration using a Qubit fluorometer (ThermoFisher Scientific, Waltham, MA) and quality using the 2100 Bioanalyzer (Agilent Technologies, Santa Clara, CA). Up to one µg of total RNA per sample was used for poly-A mRNA selection, then cDNA was synthesized from enriched and fragmented RNA using SuperScript II reverse transcriptase (Invitrogen, Carlsbad, CA) and random primers. The cDNA was further converted into double stranded DNA (dsDNA), and the resulting dsDNA was enriched with PCR for library preparation. The PCR-amplified library was purified using Agencourt AMPure XP beads (Beckman Coulter, Brea, CA). The concentration of the amplified library was measured with a Qubit fluorometer and an aliquot of the library was resolved on a Bioanalyzer. Sample libraries were multiplexed and sequenced on a NovaSeq 6000 platform (Illumina) using 75bp single-end sequencing. On average, about 30 million reads were generated from each sample.

### Data analysis of mRNAs

Raw sequencing data was demultiplexed and converted to fastq format by using *bcl2fastq* v2.20 (Illumina, San Diego, California). Then reads were aligned to the transcriptome using STAR v2.6.1 (19) RSEM v1.2.28 (20) with default parameters, using a custom human GRCh38 transcriptome reference downloaded from http://www.gencodegenes.org, containing all protein coding and long non-coding RNA genes based on human GENCODE version 23 annotation. Principal components analysis (PCA) was performed with *DESeq2* Bioconductor package v1.36.0 using expression counts for each gene in all samples, normalized for sequencing depth, and transformed with a variance stabilizing transformation. The 500 most variable genes were input into R function *prcomp,* plotted with *ggplot2* v.3.3.6, and labeled with *ggrepel* 0.9.1.

### Post-sequencing quality control and sample elimination

Initial mRNA-sequencing included 142 first trimester and 50 third trimester placenta samples, but PCA analysis showed outliers. Diagnostic heatmaps were created with R package *heatmaply* v1.1.1 to identify traces of maternal decidua contamination using genes from our sex differences analysis of first trimester placenta (NCBI GEO: GSE109082) (16) and decidua-upregulated genes (FDR<10^-20^, FC>25, FPKM>5) from our matched placenta versus decidua RNA-seq (NCBI GEO: GSE131874) (21). Samples were removed if they had traces of decidua contamination, high chromosome Y signal in presumed 46,XX samples, or other outlier status. The final “full cohort” for analysis consisted of 124 first trimester and 43 third trimester samples. PCA plots confirmed separation of first versus third trimester in the first component.

### Differential expression model

Each gene was fitted into a negative binomial generalized linear model adjusted for fetal sex, and the Wald test was applied to assess the differential expressions between first versus third trimester placenta by *DESeq2* (22, 23). The Benjamini-Hochberg False Discovery Rate (FDR) procedure was applied to adjust P-values for multiple hypothesis testing. Differential expression analysis was performed for the full cohort (124 first versus 43 third trimester samples) and for a subanalysis of 46 matched samples (23 subjects).

### Selection of thresholds

Expressed genes were defined “above sequencing noise” as determined by Y-linked gene signal from 46,XX female placenta, excluding pseudoautosomal regions. The maximum TPM from female samples was 4.01 on *PCDH11Y*, an outlier 6 orders of magnitude higher than the next gene, *RPS4Y1* (maximum TPM=0.66). Although *PCDH11Y* is not in the canonical pseudoautosomal regions of chromosome Y, it is located in a recent human-specific transposition from Xq21.3 which retains very high X/Y homology (24). Therefore, *PCDH11Y* was ignored along with pseudoautosomal genes and an expression threshold of mean TPM>0.66 was selected. Differentially expressed genes (DEGs) were defined using initial thresholds of FDR<0.05 and mean TPM>0.66 in either trimester. “Strict DEGs” were defined after the full cohort analysis (167 samples) and matched subanalysis (46 samples) were compared, leaving a robust set of “strict DEGs” with FDR<0.001, absolute fold change (FC) >1.5, and mean TPM>0.66 in either trimester.

Stably expressed genes (SEGs) were defined as genes with expression at both trimesters, excluding genes with possible differential expression and and/or high variability. Specifically, thresholds for SEGs were mean TPM>0.66 in both groups, unadjusted P-value>0.05, absolute FC<1.5, and coefficient of variation<0.80 (calculated as standard deviation divided by mean TPM from combined first and third trimester sample data).

### Heatmap

For visualization of gene expression in pathways of interest, a matrix of log_2_(TPM) values for each sample was scaled and centered, then used to create a heatmap and dendrogram, with gene rows sorted by fold change and sample columns hierarchically clustered. Figures were created with R packages *gplots* v3.1.1 and *ggplot2* v3.3.6, using function *heatmap.2*.

### Pathway enrichment analysis

Genes of interest were analyzed for enrichment in canonical pathways using Ingenuity Pathway Analysis (IPA) software (QIAGEN, Redwood City, CA, USA, www.qiagenbioinformatics.com/IPA) as previously described (16, 25). Statistical output included P-values (Fisher’s exact test) and z-scores of predicted pathway activation or inactivation. QIAGEN suggests a threshold of z-score>2 either direction for significant activation or inactivation, with less reliable predictions if z-scores are 2 to -2.

## Results

### Cohort demographics

Of the 124 first trimester and 43 third trimester placenta samples analyzed, 23 subjects overlapped. The demographics of first versus third trimester cohorts were compared without these 23 overlapping subjects (**Table 1**). There were significantly more Caucasian parents and fetuses in the first trimester only group. Race and ethnicity did not segregate in principal components analyses (PCA) of the mRNA transcriptome (**Figure S1**). Maternal pre-pregnancy BMI (21.7 versus 24.0, p=0.03) and thyroid disorders requiring thyroid replacement (3 versus 6, or 2.97% versus 30%, p=0.001) were significantly greater in the third trimester only group. There were 6 pregnant women (30%) who developed hypertension in the third trimester only group (5 required magnesium), compared to none in the first trimester placenta only group (p<0.001) (**Table 1**).

**Table 1.**
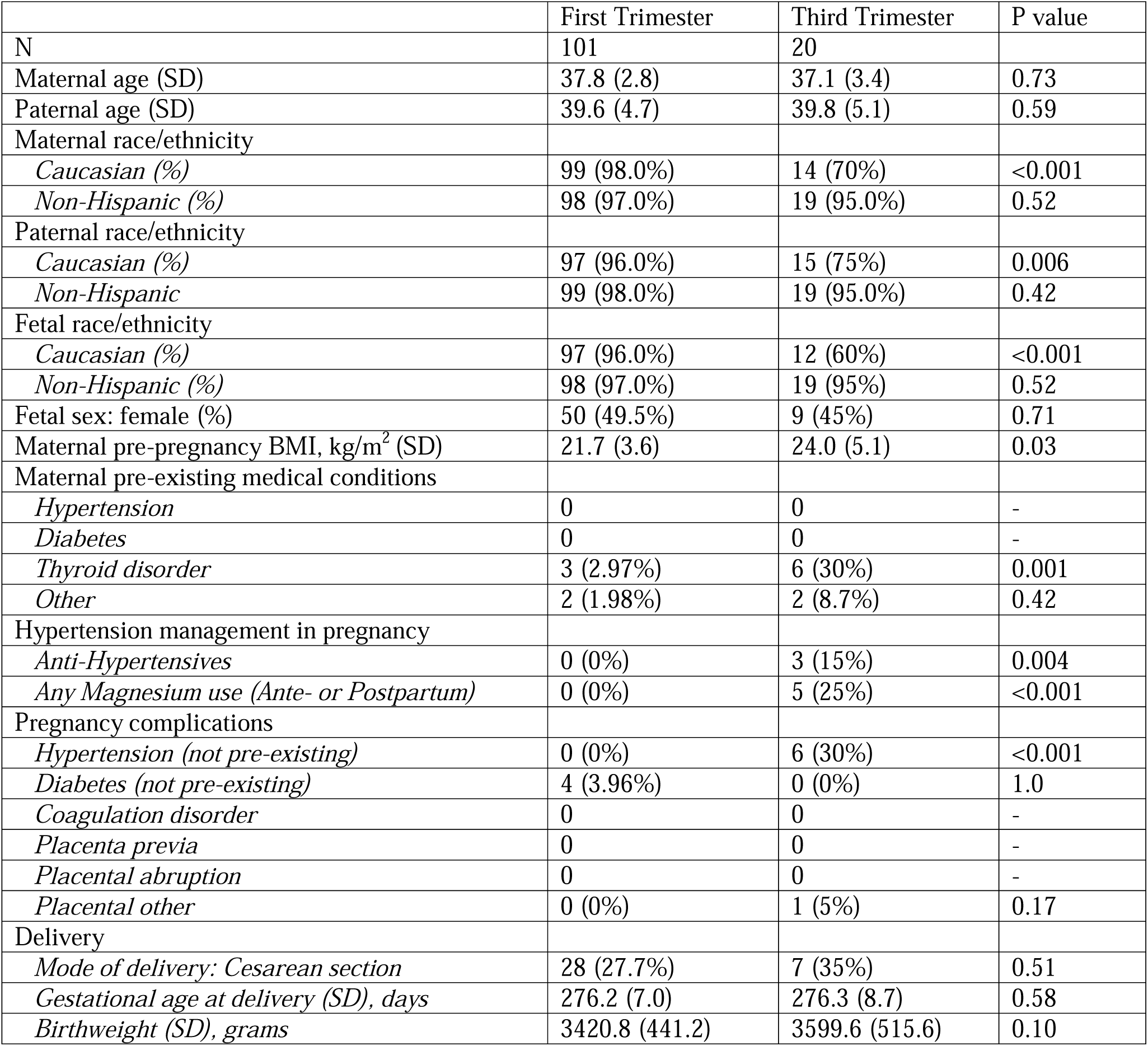
Demographics and Outcomes, Without Overlapping Pregnancies. *Table 1 caption: Numbers are formatted as “mean (standard deviation)” or “count (percentage)”. This table does not include the 23 subjects sampled at both trimesters, but they are included as a third group in supplemental demographics. SD = standard deviation.

|  | First Trimester | Third Trimester | P value |
| --- | --- | --- | --- |
| N | 101 | 20 |  |
| Maternal age (SD) | 37.8 (2.8) | 37.1 (3.4) | 0.73 |
| Paternal age (SD) | 39.6 (4.7) | 39.8 (5.1) | 0.59 |
| Maternal race/ethnicity |  |  |  |
| *Caucasian (%)* | 99 (98.0%) | 14 (70%) | <0.001 |
| *Non-Hispanic (%)* | 98 (97.0%) | 19 (95.0%) | 0.52 |
| Paternal race/ethnicity |  |  |  |
| *Caucasian (%)* | 97 (96.0%) | 15 (75%) | 0.006 |
| *Non-Hispanic* | 99 (98.0%) | 19 (95.0%) | 0.42 |
| Fetal race/ethnicity |  |  |  |
| *Caucasian (%)* | 97 (96.0%) | 12 (60%) | <0.001 |
| *Non-Hispanic (%)* | 98 (97.0%) | 19 (95%) | 0.52 |
| Fetal sex: female (%) | 50 (49.5%) | 9 (45%) | 0.71 |
| Maternal pre-pregnancy BMI, kg/m<sup>2</sup> (SD) | 21.7 (3.6) | 24.0 (5.1) | 0.03 |
| Maternal pre-existing medical conditions |  |  |  |
| *Hypertension* | 0 | 0 | - |
| *Diabetes* | 0 | 0 | - |
| *Thyroid disorder* | 3 (2.97%) | 6 (30%) | 0.001 |
| *Other* | 2 (1.98%) | 2 (8.7%) | 0.42 |
| Hypertension management in pregnancy |  |  |  |
| *Anti-Hypertensives* | 0 (0%) | 3 (15%) | 0.004 |
| *Any Magnesium use (Ante- or Postpartum)* | 0 (0%) | 5 (25%) | <0.001 |
| Pregnancy complications |  |  |  |
| *Hypertension (not pre-existing)* | 0 (0%) | 6 (30%) | <0.001 |
| *Diabetes (not pre-existing)* | 4 (3.96%) | 0 (0%) | 1.0 |
| *Coagulation disorder* | 0 | 0 | - |
| *Placenta previa* | 0 | 0 | - |
| *Placental abruption* | 0 | 0 | - |
| *Placental other* | 0 (0%) | 1 (5%) | 0.17 |
| Delivery |  |  |  |
| *Mode of delivery: Cesarean section* | 28 (27.7%) | 7 (35%) | 0.51 |
| *Gestational age at delivery (SD), days* | 276.2 (7.0) | 276.3 (8.7) | 0.58 |
| *Birthweight (SD), grams* | 3420.8 (441.2) | 3599.6 (515.6) | 0.10 |

The 23 subjects with matched first and third trimester placenta were included as a third group for the purposes of demographics analysis (**Table S1**). Compared to the first or third trimester only cohorts, paternal age was lower in the matched group. The percentage of Caucasian mothers in the matched group was lower than in the first trimester only cohort. The percentage of Hispanic parents was higher than both cohorts. The percentage of Caucasian fetuses was lower than in the first trimester only cohort. There were no additional cases of thyroid disorders requiring medication. There were 3 (13%) pregnant women in the matched group who developed hypertension, which was greater than the first trimester only cohort; one required anti- hypertensive medication and 2 required magnesium sulfate prophylaxis.

### All expressed genes (AEGs) in first and third trimester placenta

We analyzed high-throughput mRNA-sequencing of 167 placenta samples (**Figure 1A**) and observed clustering by trimester on PCA plots (**Figure 1B**, **Figure S1)**. Sequencing identified 25,312 polyadenylated RNAs in human placenta, with 14,979 RNAs expressed above sequencing noise at TPM>0.66 (**Figure S2, Figure S3**). Protein coding genes were more highly expressed than long noncoding genes, with third trimester showing higher maximum expression in both biotypes (**Figure 1C**). The most highly expressed protein coding genes in both first and third trimesters were the placental lactogen encoding genes, *CSH1* and *CSH2*, however their expression was an order of magnitude higher in third (TPM 182,551 and 82,060 respectively) compared to first trimester (TPM 17,936 and 13,339) and other expressed genes overall (**Figure 1D**). Expression differences were evident but less drastic among the next most highly expressed genes: *KISS1, CGA, EEF1A1, TFPI2, ADAM12, HBG2, TPT1* (**Figure 1D**). A shift in predominant gene families across gestation was seen in Chromosome 19, with higher expression in first trimester placenta of the chorionic gonadotropin subunits (*CGB5, CGB8, CGB*), and higher expression in the third trimester placenta of pregnancy specific beta-1-glycoprotein genes (*PSG4, PSG1, PSG3*). *PAGE4* on chromosome X is highly expressed in first trimester with mean TPM=4,517, then reduced in third trimester to mean TPM=880 (**Figure 1D**). Overall counts of placenta expressed genes were similar between first and third trimester. Placenta expressed 14,132 protein coding genes total from either trimester, with 13,664 genes in first trimester and 13,526 in third trimester (**Figure 1E**). Expressed mRNAs derived from all 22 autosomes, X and Y chromosomes, and the mitochondrial chromosome (**Figure 1D**).

**Figure 1.**
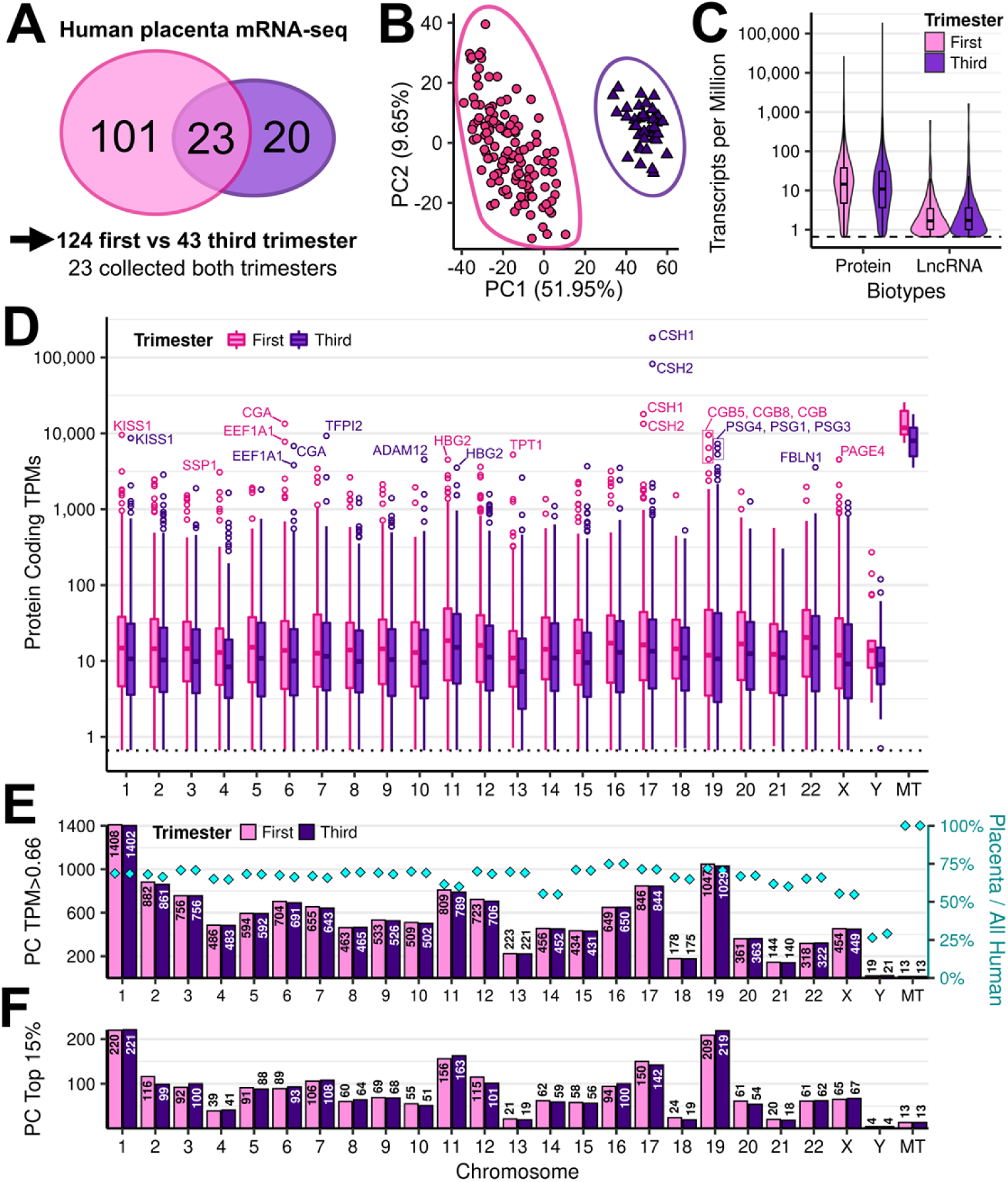
Expressed mRNAs in first and third trimester placenta. (A) The full cohort includes 124 first trimester and 43 third trimester placenta samples. (B) Principal components analysis shows separation by trimester with PC1. (C) Violin plots show the expression ranges of protein coding (Protein) and long noncoding RNAs (LncRNA). Box and whisker plots are super-imposed. The dotted line indicates the study-specific expression threshold of TPM=0.66. Only genes TPM>0.66 in either trimester are used in this figure. (D) Box and whisker plots of protein coding gene log_10_(TPM) values from first trimester (pink) and third trimester (purple) placenta, per chromosome. Points are outliers more than 1.5 times outside the interquartile range. (E) Bar plot of all expressed protein coding genes at each trimester, per chromosome. The diamond points and right y-axis indicate the proportion expressed out of all human protein coding genes, per chromosome. (F) Bar plot of the top 15% of protein coding genes, per chromosome.

The highest numbers of expressed protein coding genes came from chromosome 1 (1408 first; 1402 third trimester) and chromosome 19 (1047; 1029), the longest and densest chromosomes respectively (26). First and third trimester showed similar percentages of expressed genes out of total encoded, per chromosome (**Figure 1E**). Chromosomes 14 expressed the lowest proportion among autosome chromosomes (55.3%; 54.9%), similar to chromosome X (55.4%; 54.8%). Chromosome Y, only present in half the subjects, had an expected low percent expressed (26.4%; 29.2%). All 13 mitochondria-encoded protein coding genes were expressed in placenta (**Figure 1E**). Among the top 15% most highly expressed genes, similar patterns were observed (**Figure 1F**).

Pathway enrichment analysis was performed to investigate the biological significance of the top 15% expressed genes in first and third trimester placenta. The most significant three pathways were the same for both trimesters: EIF2 signaling (P=3.98×10^-74^ first trimester; P=1.26×10^-73^ third trimester), mitochondrial dysfunction (P=2.51×10^-68^; P=7.94×10^-48^), and oxidation phosphorylation (P=1.26×10^-63^; P=2.51×10^-42^) (**Figure S4**). Following the 3 top pathways, the next most significant pathways varied in order between first trimester and third trimester, but results were overall similar (**Table S2A, Table S2B**).

### Initial differential expression analysis demonstrated vast changes by gestational age

We performed differential expression analysis on the full cohort between first and third trimester placenta (adjusted for fetal sex) and found that the majority of polyadenylated RNAs significantly changed throughout gestation, with 12,986 differentially expressed genes (DEGs) at FDR<0.05 and TPM>0.66, representing 86.7% of all 14,979 expressed genes (TPM>0.66) in placenta (**Table S3A**). This proportion of DEGs out of all expressed genes was consistent for protein coding (86.7%; 12,253/14,132) and long noncoding genes (86.5%; 733/847). To increase confidence that these changes were due to gestational age differences, we performed a subanalysis using the 23 subjects with matched placenta samples collected at first and third trimester (**Table S3B**). This subanalysis removes the variables of genetic, demographic, and environmental differences. The subanalysis identified 11,608 DEGs (FDR<0.05, TPM>0.66), representing 77.0% of 15,071 expressed genes (TPM>0.66) in the matched cohort (**Table S3B**). The subanalysis ratios of DEGs/expressed genes were high for both protein coding genes (77.3%; 10,968/14,191) and polyadenylated long noncoding genes (72.7%; 640/880), showing that most placental mRNAs are also significantly changed from first to third trimester in a subanalysis controlled for external factors.

### Stricter thresholds identify the most robust changes across gestation

Subsequently, protein coding DEGs (FDR<0.05, TPM>0.66) from the full cohort analysis and matched subjects subanalysis were compared for congruency to identify strict DEG thresholds that serve to control for any genetic, demographic, and environmental factors (**Table S3C**). Out of 12,850 total pooled DEGs, most were significantly upregulated in the same direction in both analyses (10,363 genes; 80.6%) or trended in the same direction but only reached significance in one analysis (2,047 genes; 15.9%). There were 440 genes (3.4%) that changed direction in the subset analysis compared to the full cohort analysis, though only 8 of these mismatched genes were significant at FDR<0.05 in both analyses (only 0.062%), and all were at lower fold changes (**S6C File**). Of these eight, the highest expressed gene was *RHOC* (TPM=276 full cohort, TPM=308 matched cohort), though fold-change was very low (FC=1.06 third/first full cohort, FC=1.12 first/third matched cohort). All direction-switching genes were FC<1.16 except *SLPI* (FC=3.16 first/third full cohort, FC=7.38 third/first matched cohort) but *SLPI* had low expression (TPM=6 full cohort, TPM=1 matched cohort). A linear model shows that the full cohort and subanalysis log_2_FC values are highly correlated, with a Pearson correlation coefficient of *Rho*=0.978 and P-value=2.2×10^-16^ (**Figure S5A**). To select genes with more robust differences between first and third trimester placenta, we filtered to “strict DEGs” using thresholds of FDR<0.001, FC>1.5, and TPM>0.66 (either trimester) which resulted in a small increase in correlation (*Rho*=0.982, P-value=2.2×10^-16^, Pearson correlation) (**Figure S5B**). There were total 7,673 strict protein coding DEGs significant in at least one of the two analyses, with 3,680 (48.0%) upregulated in first trimester and 3,977 (51.8%) upregulated in third trimester placenta, and 16 (0.2%) of inconsistent direction only significant in one analysis, and none of inconsistent direction significant in both analyses. Box and whisker plots for each analysis show more genes upregulated in third trimester than first trimester with either set of thresholds but this was more pronounced with the stricter threshold (**Figure S5**). All subsequent analyses were performed with protein coding genes and the strict DEGs in the full cohort analysis, resulting in 6,941 DEGs or 49.1% of all expressed protein coding genes.

### Stably expressed genes (SEGs) throughout gestation

We identify stably expressed genes (SEGs) in placenta across gestation, meaning genes which are expressed in both first and third trimester placenta, not statistically different, and less variable between subjects. Scatter plots of the coefficients of variation for the full cohort analysis versus matched subset analysis were color-coded by different variables. High P-values did not correlate with low variability, demonstrating that high P-value alone is insufficient to define similar expression between trimesters (**Figure 2A**). However, low variability genes did trend towards low absolute FC values (**Figure 2B**). Overall expression of each gene was highly correlated between analyses, with a Pearson coefficient of *Rho*=0.997 and linear model P-value=2.2×10^-16^ for mean TPM per trimester (**Figure 2C**). Based on these analyses, thresholds were selected to remove highly variable genes and better define SEGs across gestation in the full cohort: TPM>0.66 in both trimesters, P-value>0.05, FC<1.5, and a coefficient of variation<0.80. Of all expressed protein coding genes, 10.9% are SEGs (1,545/14,132).

**Figure 2.**
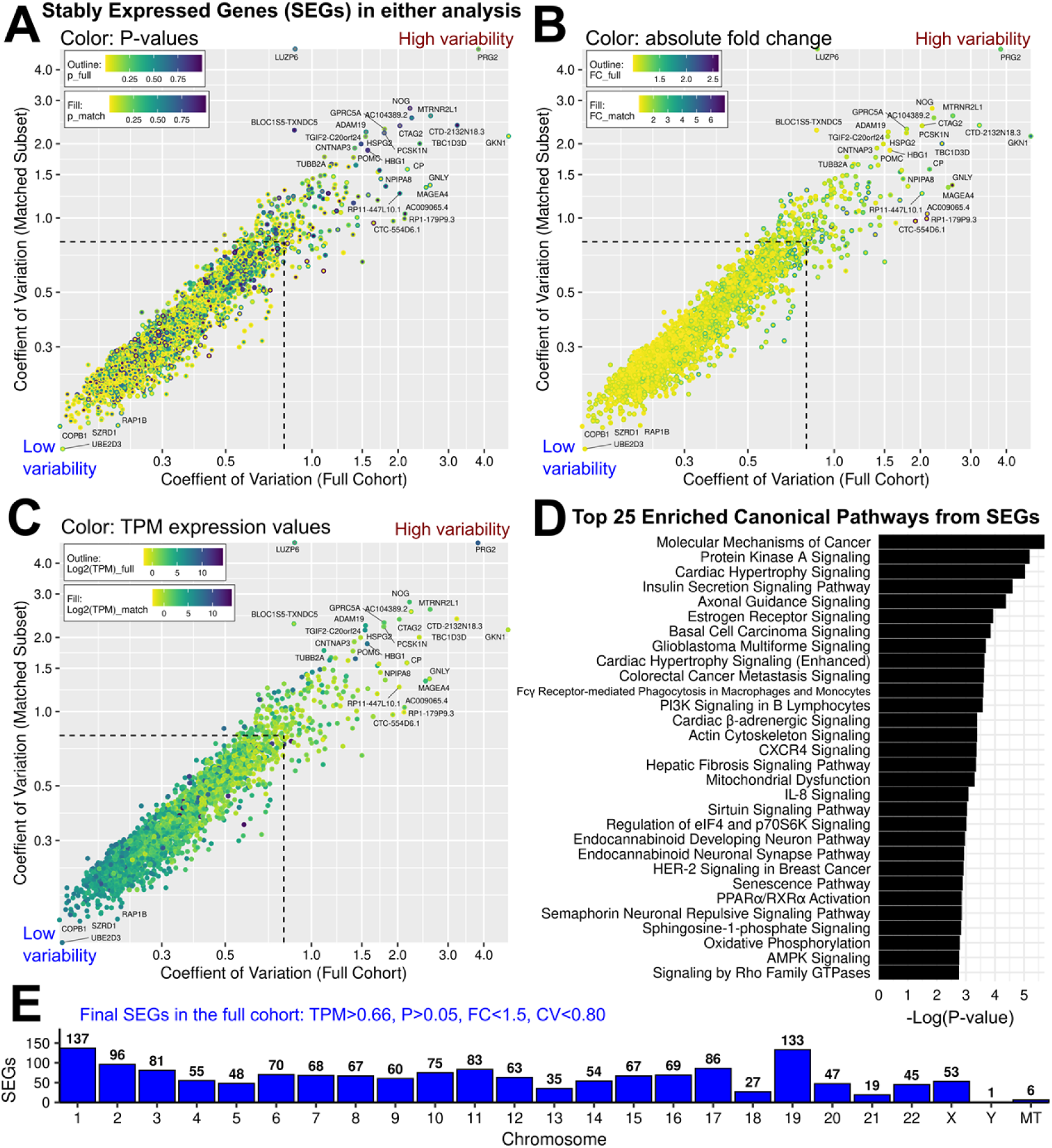
Variability and selection of stably expressed genes. Comparison of the full cohort analysis (124 first versus 43 third trimester samples) with a subanalysis of only matched first and third trimester samples (first versus third trimester from the same 23 pregnancies). (A-C) Scatter plots comparing the coefficients of variation (CV) from each analysis, per gene. Genes are plotted if TPM>0.66 (both trimesters), P-value>0.05, and FC<1.5 in either analysis. Stably expressed genes (SEGs) are defined by those three filters and also CV<0.80. Dashed lines showCV=0.80. Points are color-coded by different variables to identify differences between the full cohort value (outline color) and subanalysis value (fill color). Axes are log2-transformed to better visualize low variability genes. Color coding based on: (A) Wald P-value, (B) fold change, (C) log_2_(TPM) expression. (D) Top 25 enriched canonical pathways among SEGs from the full cohort. (E) Bar plot of SEGs from the full cohort.

Pathway enrichment analysis of protein coding SEGs in placenta showed significant enrichment predominantly for cell growth and proliferation pathways (molecular mechanisms of cancer, basal cell carcinoma signaling, glioblastoma multiforme signaling); cell movement and cell-cell communication (protein kinase A signaling, cardiac hypertrophy signaling, insulin secretion signaling, axonal guidance signaling); hormone and small molecule signaling (insulin secretion signaling, estrogen receptor signaling) (**Figure 2D, Table S2C**).

The chromosome distributions of SEGs were similar to total mRNA expression, with most genes from chromosome 1 (137 SEGs), then chromosome 19 (133 SEGs) (**Figure 2E**). The fewest SEGs were from chromosome Y (1 SEG), the mitochondrial genome (6 SEGs), and chromosome 21 (19 SEGs) (**Figure 2E**).

### Strict DEGs in the full cohort

The full cohort analysis identified 6,941 protein coding strict DEGs (FDR<0.001, FC>1.5, TPM>0.66), composed of 3,206 upregulated in first trimester and 3,735 upregulated in third trimester placenta. The volcano plot shows more third trimester upregulated DEGs with high fold change (FC), compared to first trimester placenta (**Figure 3A**). The most highly expressed DEGs were chorionic somatomammotropin hormone 1 and 2, which encode human placental lactogen: *CSH1* with TPM increase from 17,936 to 182,551 in first to third trimester (14.0-fold, FDR 1.24×10^-40^) and *CSH2* with TPM increase from 66,113 to 165,274 in first to third trimester (8.3- fold, FDR 5.36×10^-24^) (**Figure 3B**). The median fold changes were higher in third trimester upregulated DEGs across all chromosomes, except chromosomes 12 and 18 which were higher in first trimester (**Figure 3C**). The DEGs with greatest fold increase in the first trimester includes *ANGPT4* with TPM decrease from 10.55 to 0.03 from first to third trimester (219.0-fold, FDR=5.17×10^-83^) and *MMP12* with TPM decrease from 33.7 to 0.11 from first to third trimester (186.3-fold, FDR=2.78×10^-75^) (**Figure 3C**). The four DEGs with greatest fold increase in the third trimester placenta are *IDO2* with TPM increase from 0.04 to 11.39 in first to third trimester (463.7-fold, FDR=4.31×10^-288^), *ALPP* with TPM increase from 9.39 to 2850.47 from first to third trimester (434.3-fold, FDR=4.27×10^-297^), *IL1R1* with TPM increase from 0.27 to 40.99 from first to third trimester (223.3-fold, FDR=1.19×10^-227^), and *FABP4* with TPM increase from 1.01 to 133.74 in first to third trimester (198.0-fold, FDR=0). All chromosomes were represented, with protein DEGs most represented on chromosome 1 (758 genes; 10.9%) and chromosome 19 (522 genes; 7.5%) (**Figure 3D**).

**Figure 3.**
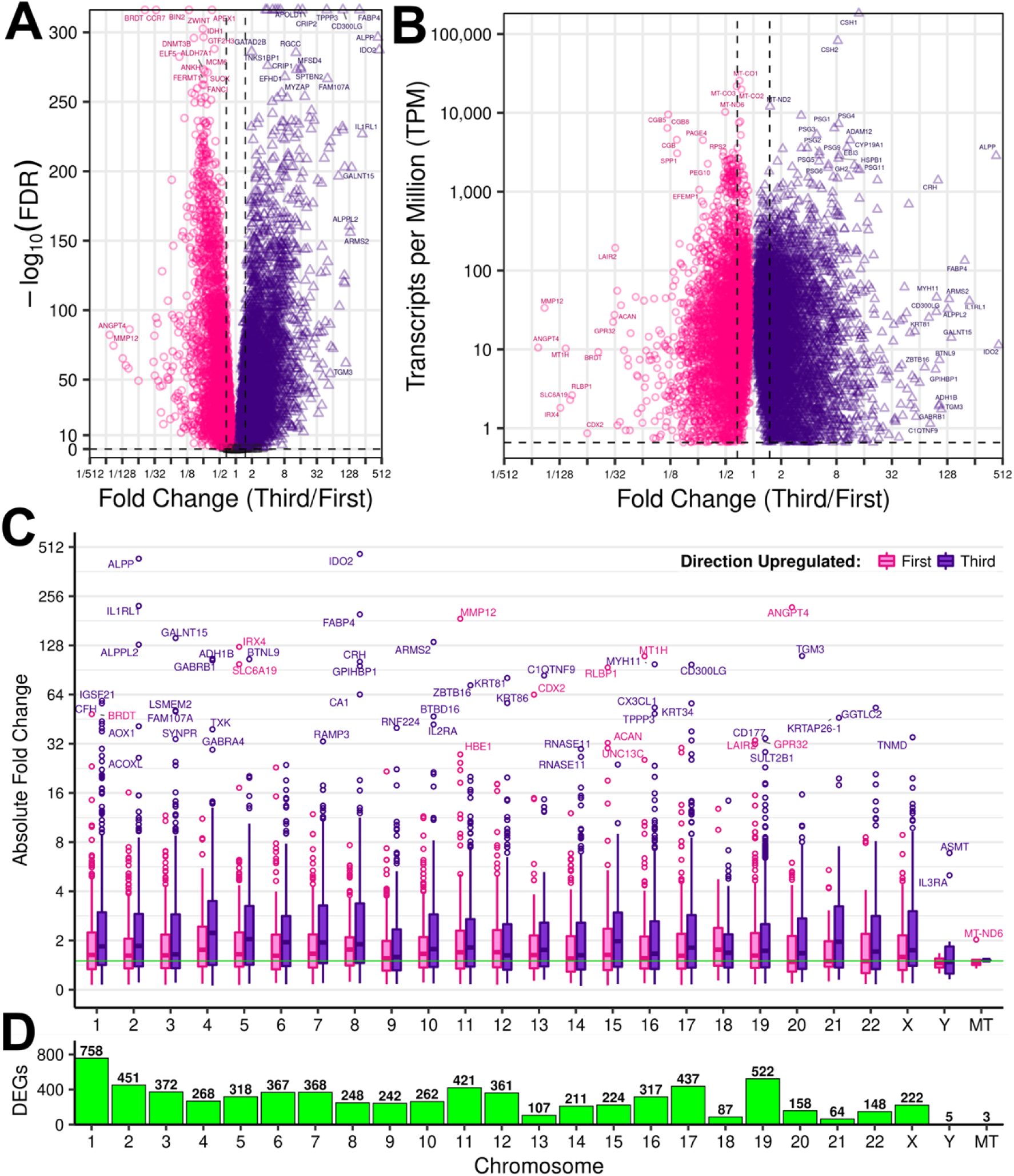
Differentially expressed protein coding genes between first and third trimester placenta (full cohort). (A) Volcano plot of all expressed protein coding genes (TPM >0.66). (B) Expression versus FC for DEGs. The dotted lines represent FC=1.5. (C) Box and whisker plots of absolute FC distribution across chromosomes for all protein coding genes at FDR<0.001 and TPM>0.66 (any FC). Solid green line is FC=1.5. (D) Chromosome frequency of all protein coding DEGs at FDR<0.001, FC>1.5, and TPM>0.66 either trimester.

Pathway enrichment analysis demonstrates first trimester upregulated DEGs were most enriched for EIF2 signaling, kinetochore metaphase signaling, cell cycle control of chromosomal replication, role of BRCA1 in DNA damage response, and superpathway of cholesterol biosynthesis (**Figure 4A, Table S2D**). Third trimester DEGs were most enriched for cardiac hypertrophy signaling (enhanced); hepatic fibrosis / hepatic stellate cell activation; role of macrophages, fibroblasts, and endothelial cells in rheumatoid arthritis; and tumor microenvironment pathway (**Figure 4B, Table S2E**). A heatmap of DEGs from either trimester in the most enriched pathway among DEGs, EIF2 signaling pathway, demonstrates sample segregation by trimester (**Figure 4C**).

**Figure 4.**
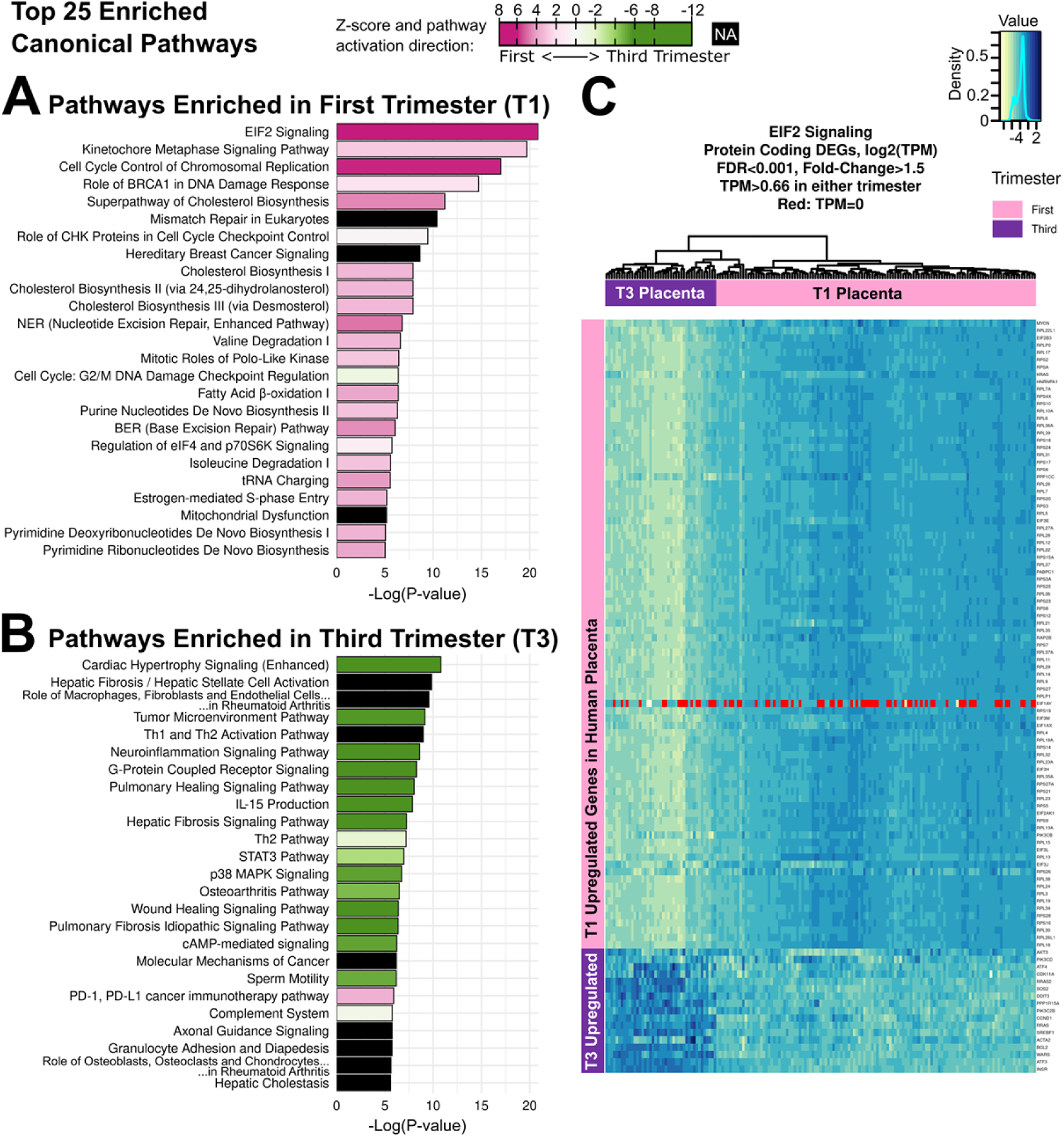
Top 25 enriched canonical pathways. Z-scores of magnitude >2 in either direction indicate significant predicted pathway activation or inactivation. Pathway predicted activation color coding: pink=activated in first trimester, green=activated in third trimester, black=not available. Enriched pathways among strict DEGs (FDR<0.001, FC>1.5, TPM>0.66 either trimester) upregulated in (A) first trimester, (B) third trimester. (C) Heatmap of log_2_(TPM) values for genes in the EIF2 signaling pathway, full cohort data. The Y-linked gene *EIF1AY* is not expressed in female samples.

## Discussion

The main component of the placenta, the chorionic villi, changes significantly over the course of gestation to meet its changing roles for the growing fetus. These differences are demonstrated by the variability of the normative placenta transcriptome across gestation, with 86.7% of expressed protein coding genes in the human placenta significantly different between first and third trimester by high-throughput transcriptomics. These differences provide insight into placental function changes across gestation. We also identified stably expressed genes (10.9% of the placental transcriptome) which may sustain core placental functions. This is the largest sample size of first and third trimester placentas of spontaneously conceived female and male singletons in ongoing and completed healthy pregnancies, constituting the normative human placenta atlas.

Chromosome distributions were consistent across all expressed, differentially expressed, and stably expressed genes. The high representation from chromosome 1 is expected as it is the largest chromosome. Chromosome 19, though much smaller, has the highest density of human genes (26). In our previous work on miRNA expression in placenta, chromosome 19 contributed the most miRNAs in both first and third trimester due to a known placenta-specific miRNA cluster unique to primates (18, 27). Together, these two studies demonstrate that chromosome 19 is critical to the human placenta epigenome and transcriptome. Chromosome 14 also contains a placenta-specific miRNA cluster and was the next greatest contributor to placenta miRNA expression, but it was relatively underrepresented in placenta protein coding gene expression compared to other chromosomes.

Chromosome distributions show that all 13 mitochondrial-encoded protein coding genes are highly expressed in placenta (among the top 15%), including six SEGs and three strict DEGs, suggesting that mitochondrial gene expression adapts to meet metabolic needs throughout gestation. The high expression of the mitochondrial genes reflects their vital role in placental utilization of oxygen to generate energy via oxidative phosphorylation (28, 29) and is consistent with the described adaptation of mitochondria to the changing oxygen demand over gestation (30). Mitochondrial pathways were two of the three most significantly enriched pathways among the top 15% placental expressed genes: “Mitochondrial Dysfunction” and “Oxidative Phosphorylation”. Mitochondrial dysfunction is implicated in oxidative stress gestational disorders including intrauterine growth restriction (IUGR), preeclampsia, gestational diabetes, and intrahepatic cholestasis of pregnancy (28, 29, 31, 32). Studies profiling the mitochondrial genes associated with these disorders are sparse. *MT-ND6* (2.03-fold upregulated in first trimester, FDR=9.67×10^-31^) as well as *MT-ATP6* (a SEG in our healthy cohort) were negatively associated with IUGR in a case control study (33). Of the four mitochondrial genes associated with change in regulation in intrahepatic cholestasis of pregnancy described in the literature (32), three were stably expressed across gestation in this study: *MT-ND4L*, *MT-CYB*, *MT-ND4*. An increased copy number of *MT-ND1* (a SEG) was found in early onset preeclampsia (<34 weeks), suggesting an adaptative response to oxidative stress (34). Further characterization of the mitochondrial biomarkers involved in oxidative stress pathologies of gestational disorders is important in aiding future clinical interventions.

The majority (86.7%) of placental mRNA transcripts significantly change from first to third trimester with FDR<0.05 alone, and 49.1% remain at our strict DEG thresholds. The two most upregulated genes in first trimester highlight the environmental challenges of early placenta development in low oxidative states: angiopoietin 4 (*ANGPT4*, 218-fold) and matrix metallopeptidase 12 (*MMP12*, 186-fold). *ANGPT4* is an angiogenic factor expressed in a variety of tissue, but there is little information on its expression in placenta. *MMP12* plays a role in low oxygen states and invasive trophoblast and may play a critical role early in placentation (35, 36). The two strict DEGs most upregulated in the third trimester include an isoform of indoleamine 2,3-dioxygenase (*IDO2*, 463-fold) and placental alkaline phosphatase (*ALPP*, 434-fold). Indoleamine 2,3-dioxygenase is an enzyme involved in tryptophan catabolism which contributes to immunotolerance of the fetus and other foreign bodies, with isoforms *IDO* and *IDO2* possibly having distinct functions in cancer and placenta (37, 38). Placental alkaline phosphatase (ALPP) is a marker of syncytiotrophoblasts at term but not first trimester (21, 39–41), and is associated with regulation of fetal growth later in pregnancy (42). Abnormal levels of ALPP have been implicated in intrauterine growth restricted pregnancies and preterm deliveries (43). The expression pattern of ALPP is well established, and this study supports other evidence that it increases throughout gestation (44, 45). Two other genes that increase dramatically in third trimester are the human placental lactogen genes (*CSH1* and *CSH2*), which had expression an order of magnitude higher than the next most highly expressed genes. Human placental lactogen promotes production of insulin-like growth factor, thus modulating fetal growth and metabolism (46, 47). The high expression of *CSH1* and *CSH2* in third trimester reflects the increasing metabolic demands of the placenta and fetus as gestation progresses.

The major strengths of this study are the use of first and third trimester tissue from healthy pregnancies that continued until delivery, the largest cohort sizes to date, and the use of high-throughput sequencing. Uniquely, this study contains chorionic villi from pregnancies sampled at both first and third trimester, which minimizes subject variability in the full cohort and reduces the effect of genetic and environmental differences, with full cohort results highly correlated to the subanalysis of only matched subjects (coefficient of 0.98). These strengths allow for a more precise evaluation of human placental transcriptomes across gestation.

The limitations of this study include some demographic differences between the first and third trimester groups, including race and ethnicity differences. However, the overall differences were minimal and PCA plots did not demonstrate clustering based on race and ethnicity. We also identified small differences in maternal BMI and thyroid disorders, but PCA plots did not show clustering or outliers. Although there was a higher rate of pregnancy complications, specifically hypertension, subjects were enrolled in the first trimester prior to development of pregnancy complications. Furthermore, the subanalysis with only overlapping subjects shows that these demographic differences have a minimal effect on results, and differences seen are driven by gestational age.

Overall, this study demonstrates that although the early and late placenta is composed of a similar structure, the chorionic villi, these villi change dramatically to support the pregnancy throughout gestation. As the largest study, and the only study with matched subjects at both collection times, this work provides a normative mRNA atlas of first and third trimester human placenta. This work serves as a reference for future studies aimed at understanding gestational age specific functional mechanisms and development of gestation specific biomarkers.

## Acknowledgements

We thank the patients who consented to participate in research, without whom none of this is possible. We thank clinical research coordinators Rae A. Buttle, Erica Sauro, Rosemarie DiPentino, Kerlly Castellano, Allynson Novoa, and Rimsha Hussaini for recruitment work. We thank the Cedars-Sinai cytogenetics staff and the midwives for help collecting samples. We thank the staff at the Cedars-Sinai Cancer Applied Genomics Shared Resource for sequencing. Research reported in this publication was supported by the National Institutes of Health under awards by the Eunice Kennedy Shriver National Institute of Child Health & Human Development (R01HD091773 to MDP), The National Institute of Allergy and Infectious Disease (R01AI154535 to MDP, R01AI164504 to CAJ), the National Institute of Biomedical Imaging and Bioengineering (U01EB026421 to MDP and HRT), and the National Heart Lung and Blood Institute (R01HL167268 to SAK). The funding agency was not involved in the design, analysis, or interpretation of the data reported. The content is solely the responsibility of the authors and does not necessarily represent the official views of the National Institutes of Health.

## Author Contributions

Conceptualization: TLG, SW, AEF, CAJ, KL, YZ, HRT, SAK, MDP

Data Curation: TLG, YW, ELC

Formal Analysis: TLG, SW, YW, JLC

Funding Acquisition: HRT, MDP

Investigation: TLG, SW, CS, ELC, NVJ, RD

Methodology: TLG, YW, CS, JLC

Project Administration: TLG, MDP

Resources: JW

Software: TLG, YW

Supervision: MDP

Visualization: TLG, YW

Writing – Original Draft Preparation: TLG, SW, MDP

Writing – Review & Editing: TLG, MDP

## Data and Code Availability

The mRNA-sequencing data has been deposited in the National Center for Biotechnology Information Gene Expression Omnibus (NCBI GEO) under accession [available upon acceptance].

