## Supplemental Information for "High-throughput mRNA-seq atlas of human placenta shows vast transcriptome remodeling from first to third trimester"

*Listed in order of appearance in the manuscript text.*

**Figure S1. Principal Components Analysis (PCA)**. PCA plots for the mRNA-seq results of the full cohort, 124 first trimester and 43 third trimester human placenta samples, 167 total. (A-J) PCA plots color coded by different variables.

**Table S1. Table of demographics and pregnancy outcomes for all sequenced subjects, with the matched cohort analyzed as a third group.** Subjects are separated into first trimester only, third trimester only, and the matched group.

**Figure S2. Quality control heatmaps.** Heatmaps to confirm expected fetal sex in this mRNA-seq data, using X/Y-linked genes significant in our previous total RNA-seq sex differences analysis (Gonzalez et al 2018, NCBI GEO:GSE109082), sorted by fold-change in that study. (A) First trimester. (B) Third trimester.

**Figure S3. Selection of the study-specific expression threshold, TPM>0.66.** Scatter plot of TPM values for Y-linked genes in females (46,XX) and males (46,XY). TPM = transcripts per million.

**Figure S4. Pathway enrichment analysis for the top 15% expressed protein coding genes in human placenta.** Bar plots of the top 25 most significant pathways. (A) First trimester. (B) Third trimester.

**Table S2. Pathway enrichment analysis result spreadsheets.** Five sheets show results for the following inputs: (A) first trimester top 15% expressed protein coding genes, (B) third trimester top 15% expressed protein coding genes, (C) SEGs, (D) strict DEGs upregulated in first trimester, (E) strict DEGs upregulated in third trimester. [SEPARATE XLSX FILE]

**Table S3. Differential expression analysis result spreadsheets.** DESeq2 differential expression analysis results from (A) the full cohort, (B) the subanalysis with 23 subjects sampled at both first and third trimester. Model: first versus third trimester, adjusted for fetal sex. (C) Comparison analysis spreadsheet with the coefficients of variation. [SEPARATE XLSX FILE]

**Figure S5. Selection of strict DEG thresholds.** (A) Scatter plot, linear regression, Pearson correlation coefficient (*R*), and P-value for all DEGs in either analysis at TPM>0.66 and FDR<0.05. Box and whisker plots are shown for each analysis in the margins. Fold change (FC) is first/third trimester, so positive $log_2$FC means upregulated in first trimester. Smaller point shapes indicate if a gene only met the expression threshold (TPM>0.66 at first and/or third trimester) in one analysis. (B) Strict DEGs are significant at TPM>0.66, FDR<0.001, and absolute FC > 1.5. Plotted as in A.

Figure S1A: Principal Components Analysis (PCA) of First and Third Trimester Human Placenta, 167 Samples

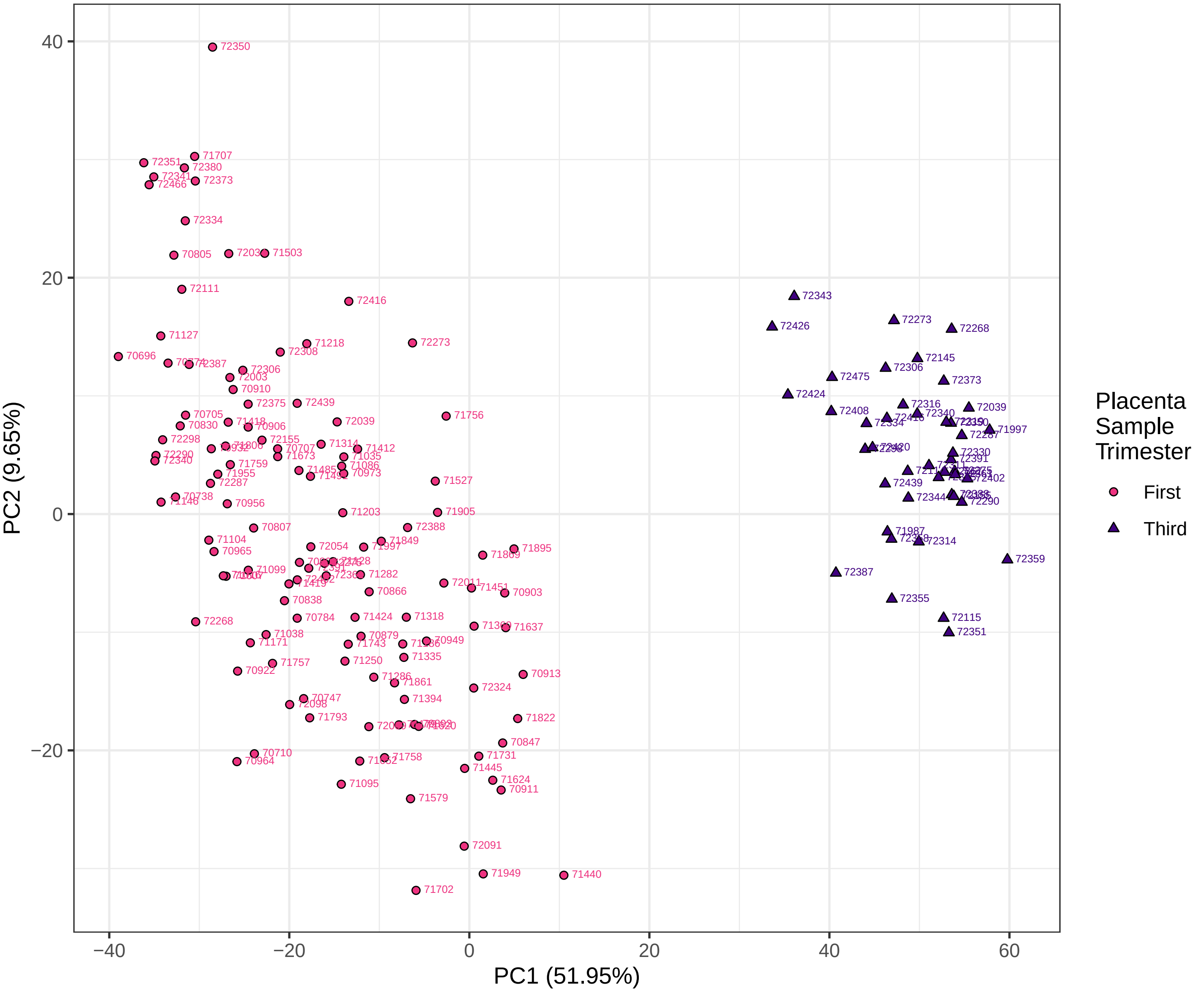

Figure S1B: PCA of First and Third Trimester Human Placenta, 167 Samples: Trimester

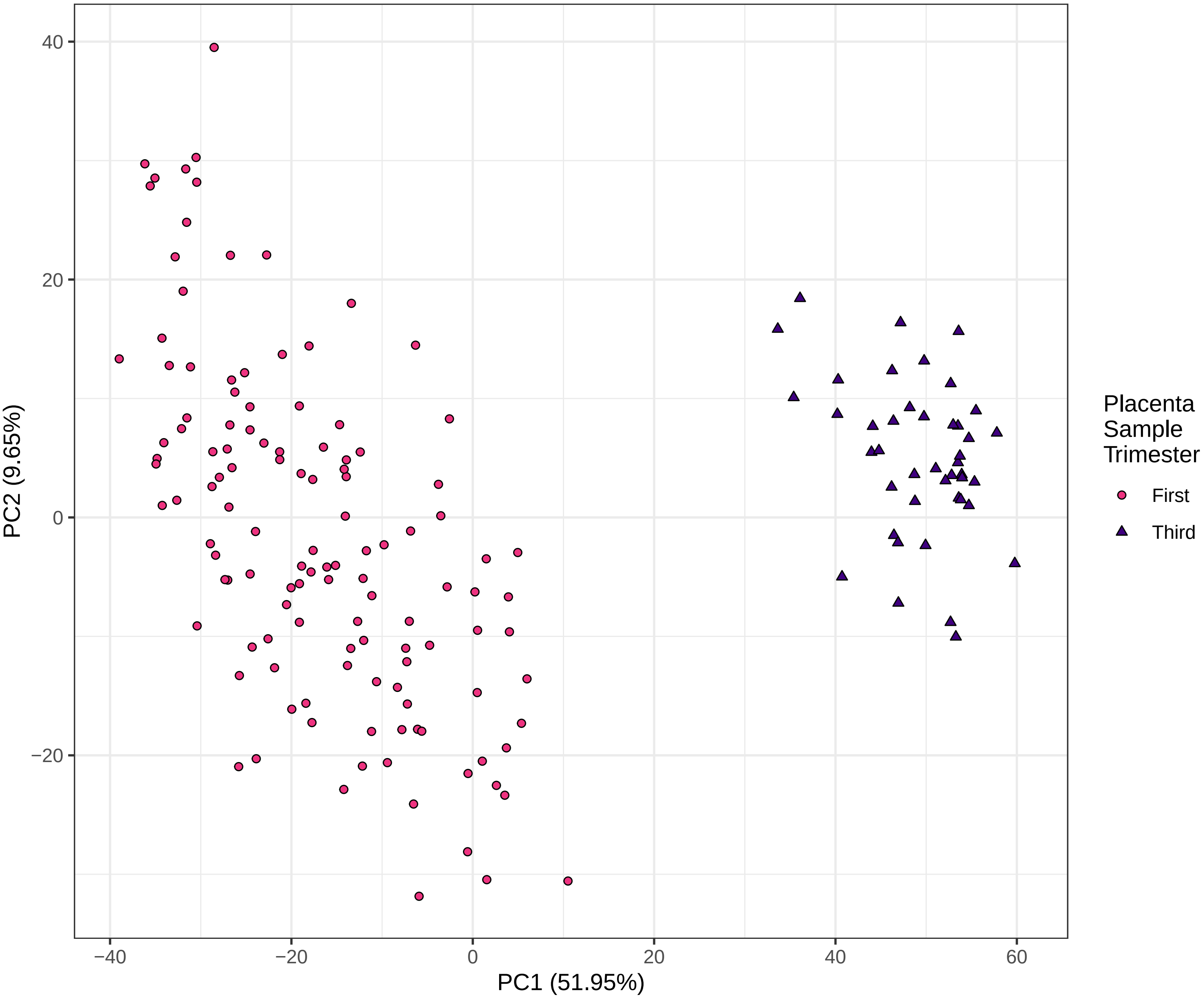

Figure S1C: PCA of First and Third Trimester Human Placenta, 167 Samples: Gestational Age at Collection

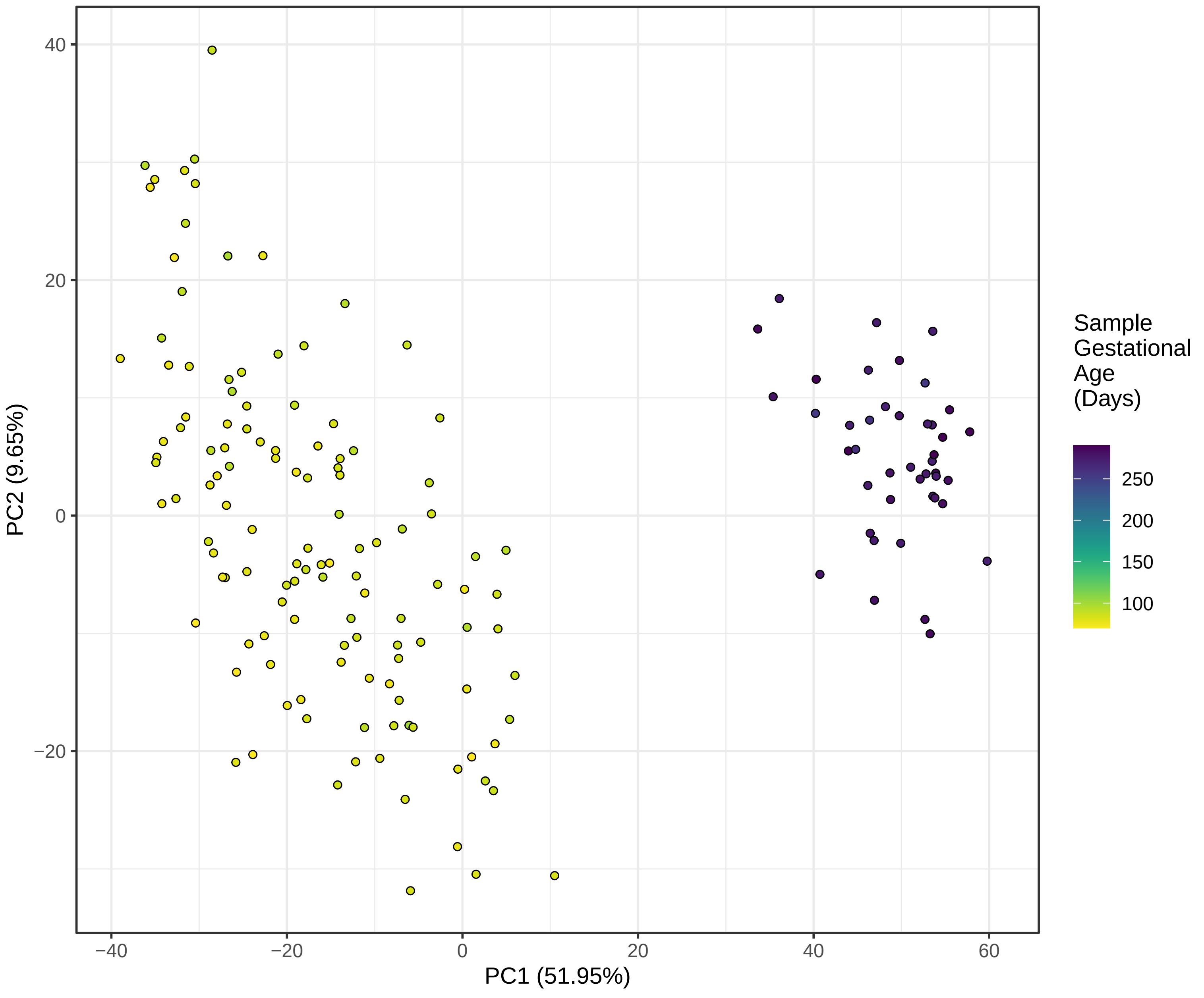

Figure S1D: PCA of First and Third Trimester Human Placenta, 167 Samples: Maternal Age

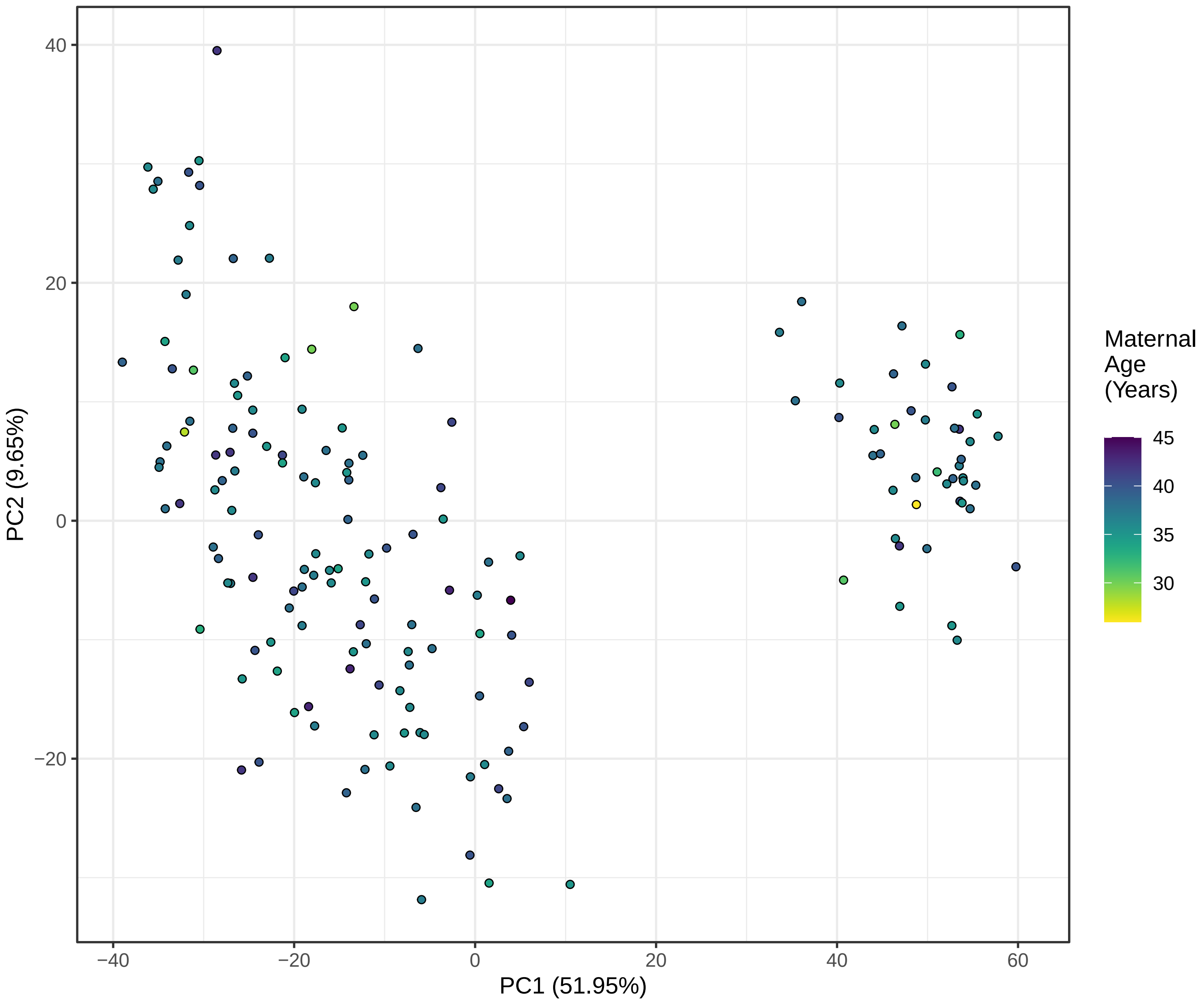

Figure S1E: PCA of First and Third Trimester Human Placenta, 167 Samples: Paternal Age

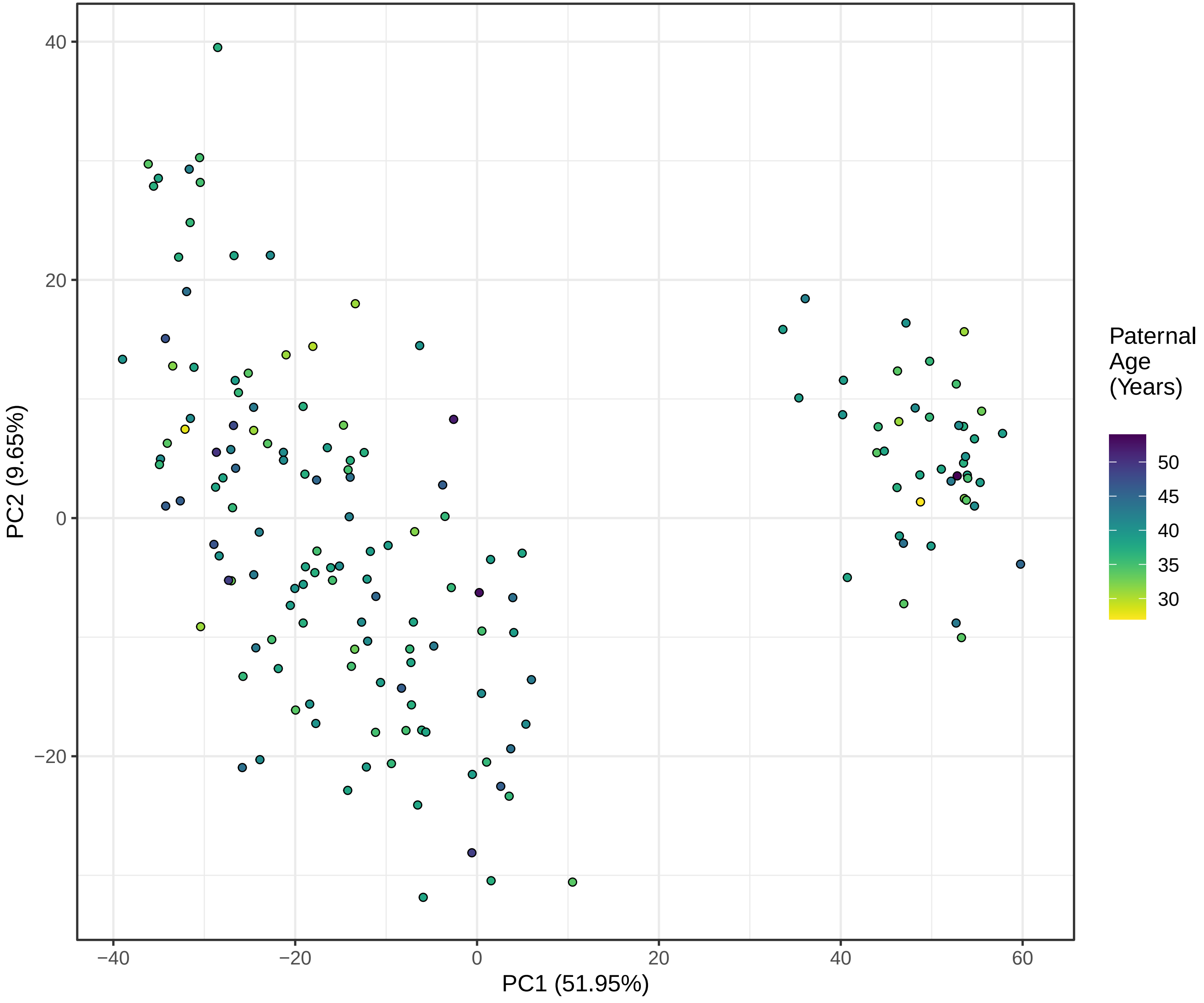

Figure S1F: PCA of First and Third Trimester Human Placenta, 167 Samples: Pre−Pregnancy Maternal BMI

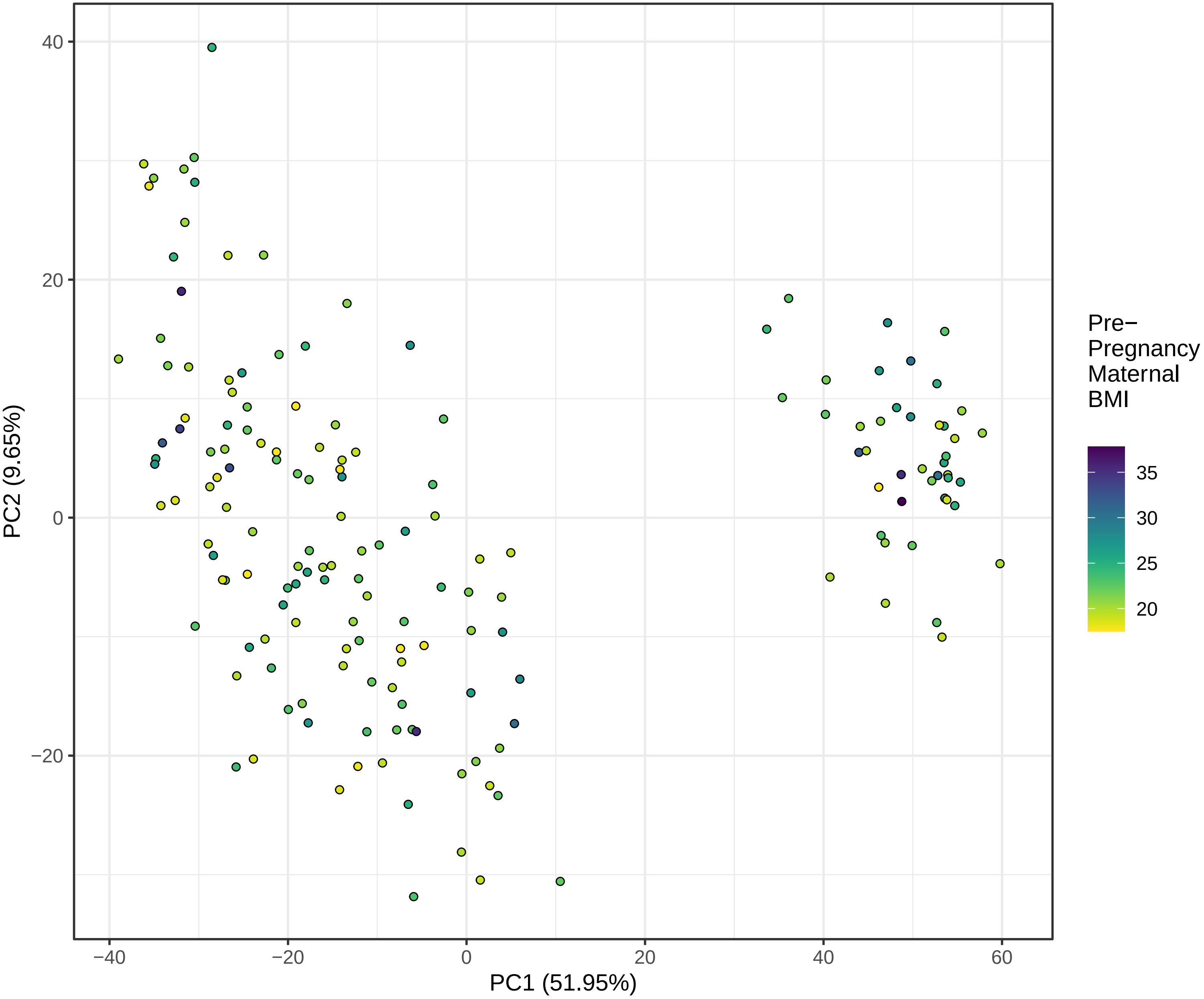

Figure S1G: PCA of First and Third Trimester Human Placenta, 167 Samples: Fetal Sex

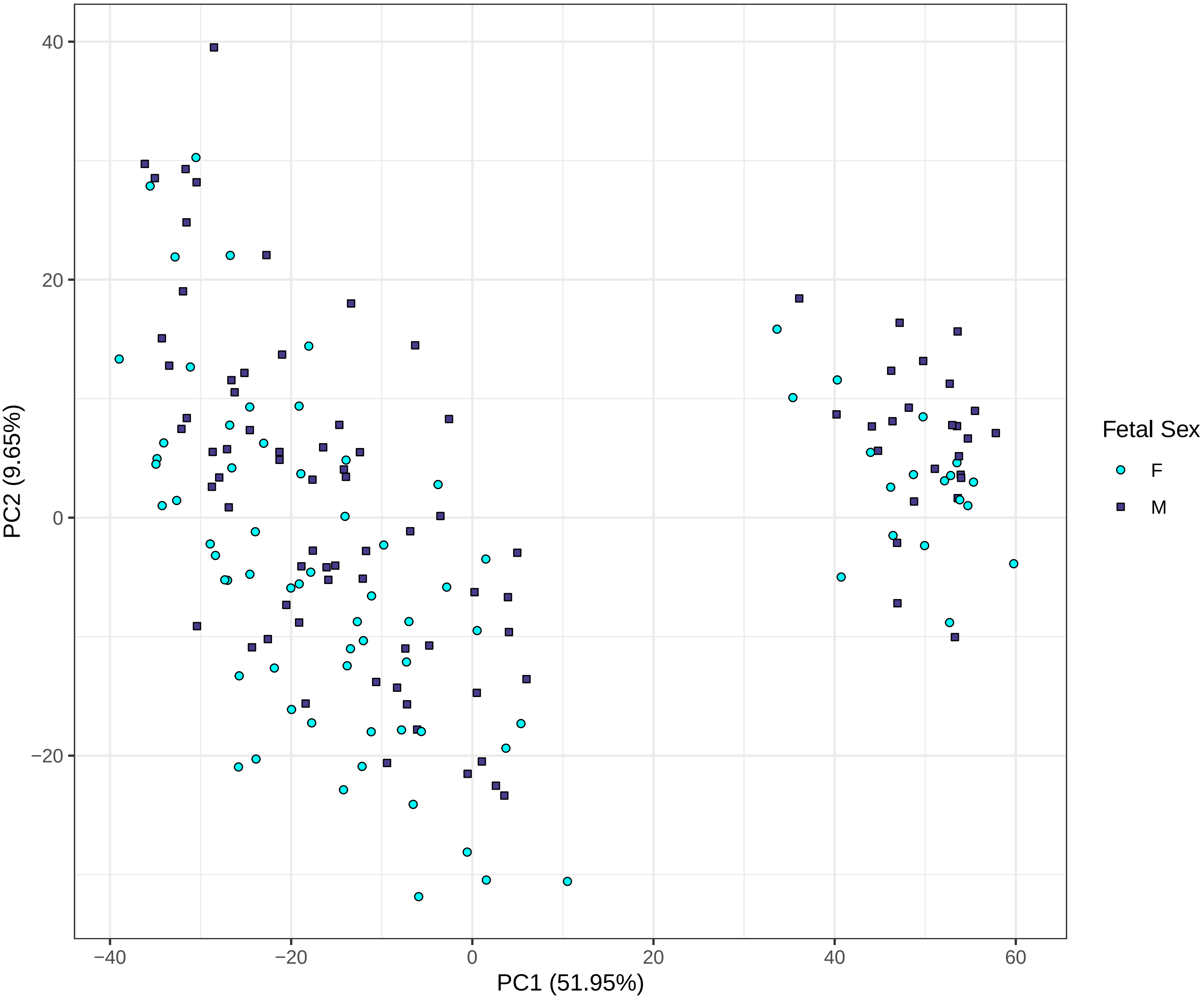

Figure S1H: PCA of First and Third Trimester Human Placenta, 167 Samples: Fetal Race

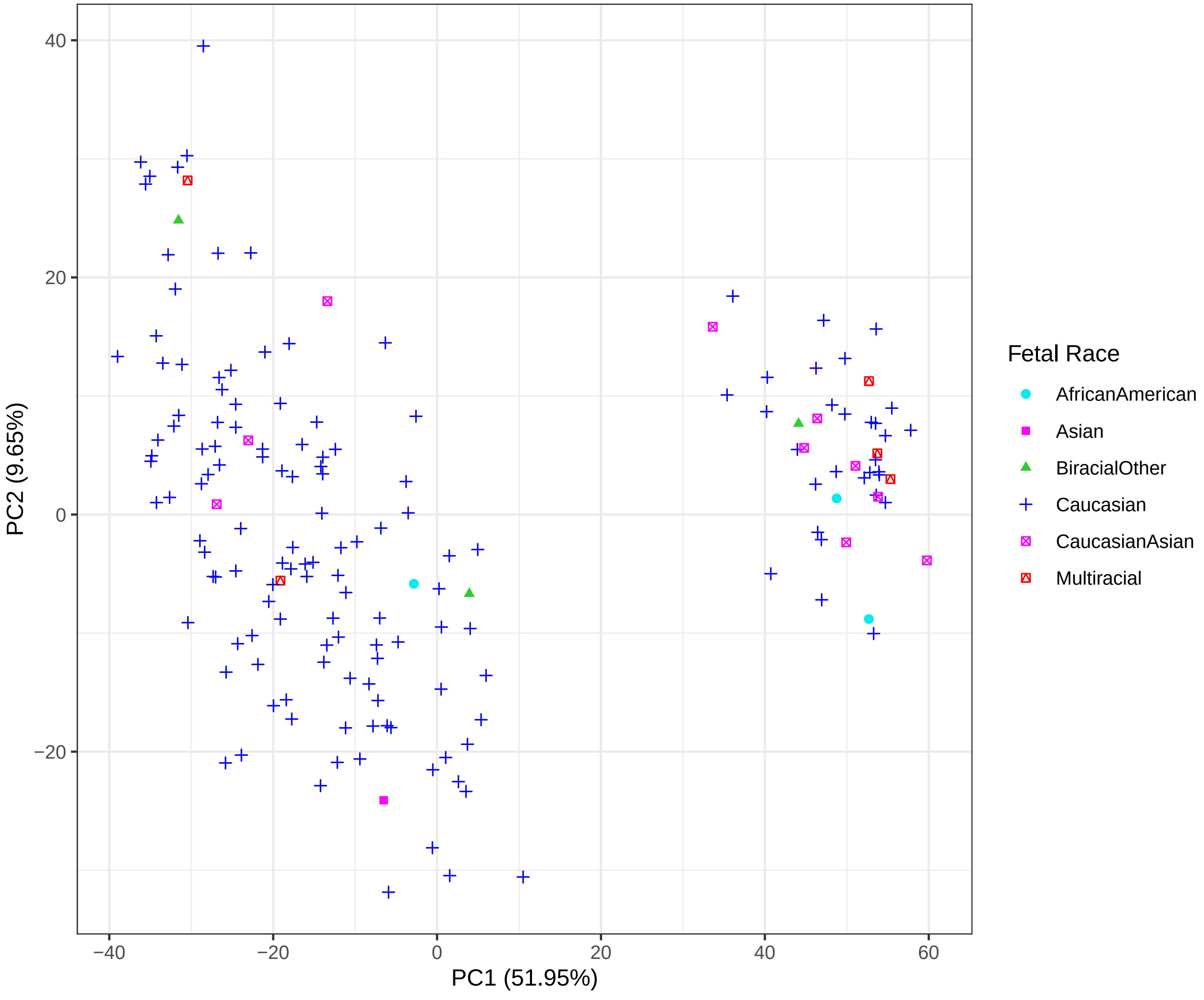

Figure S1I: PCA of First and Third Trimester Human Placenta, 167 Samples: Fetal Ethnicity

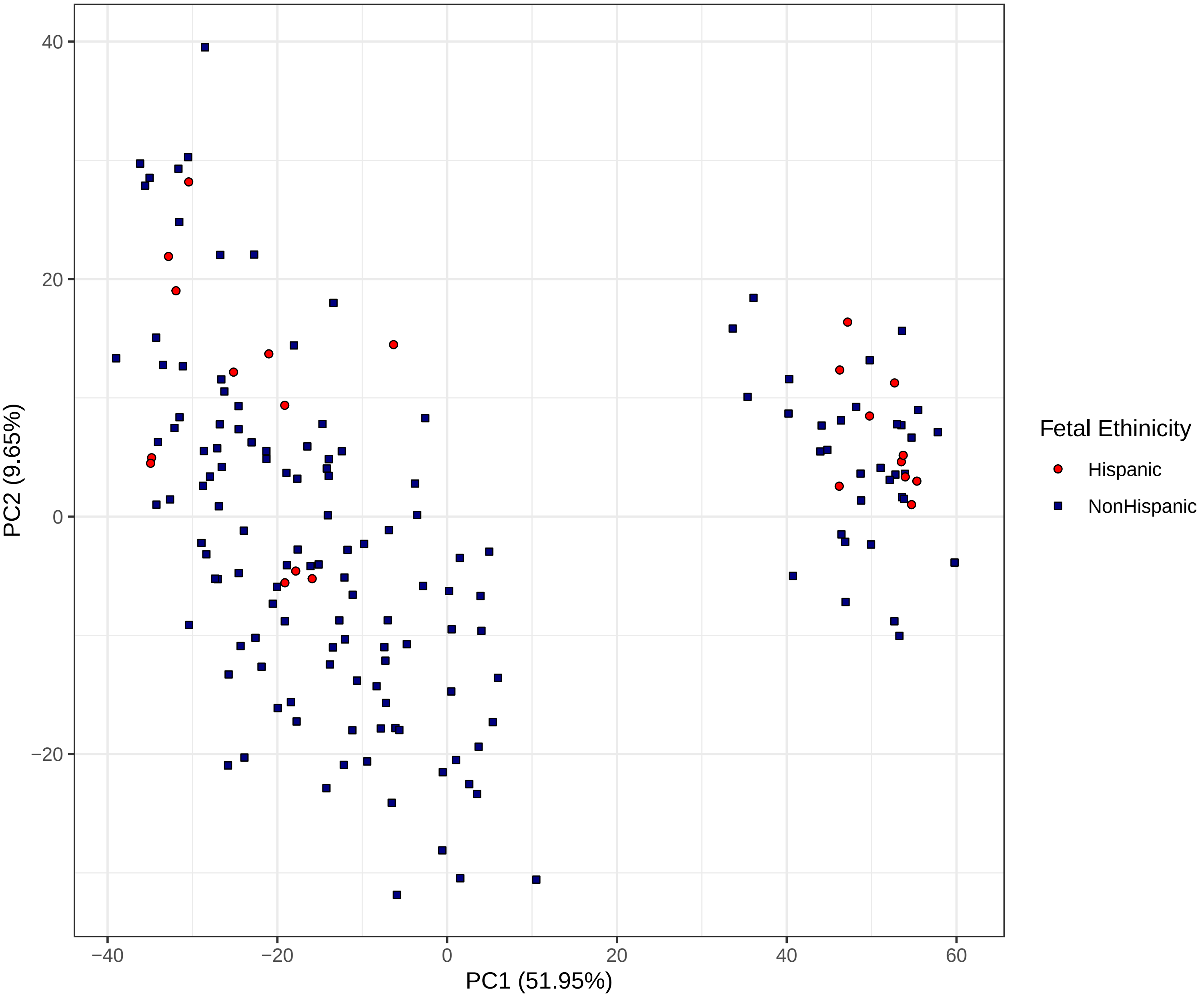

Figure S1J: PCA of First and Third Trimester Human Placenta, 167 Samples: RNA Integrity Number (RIN) Value

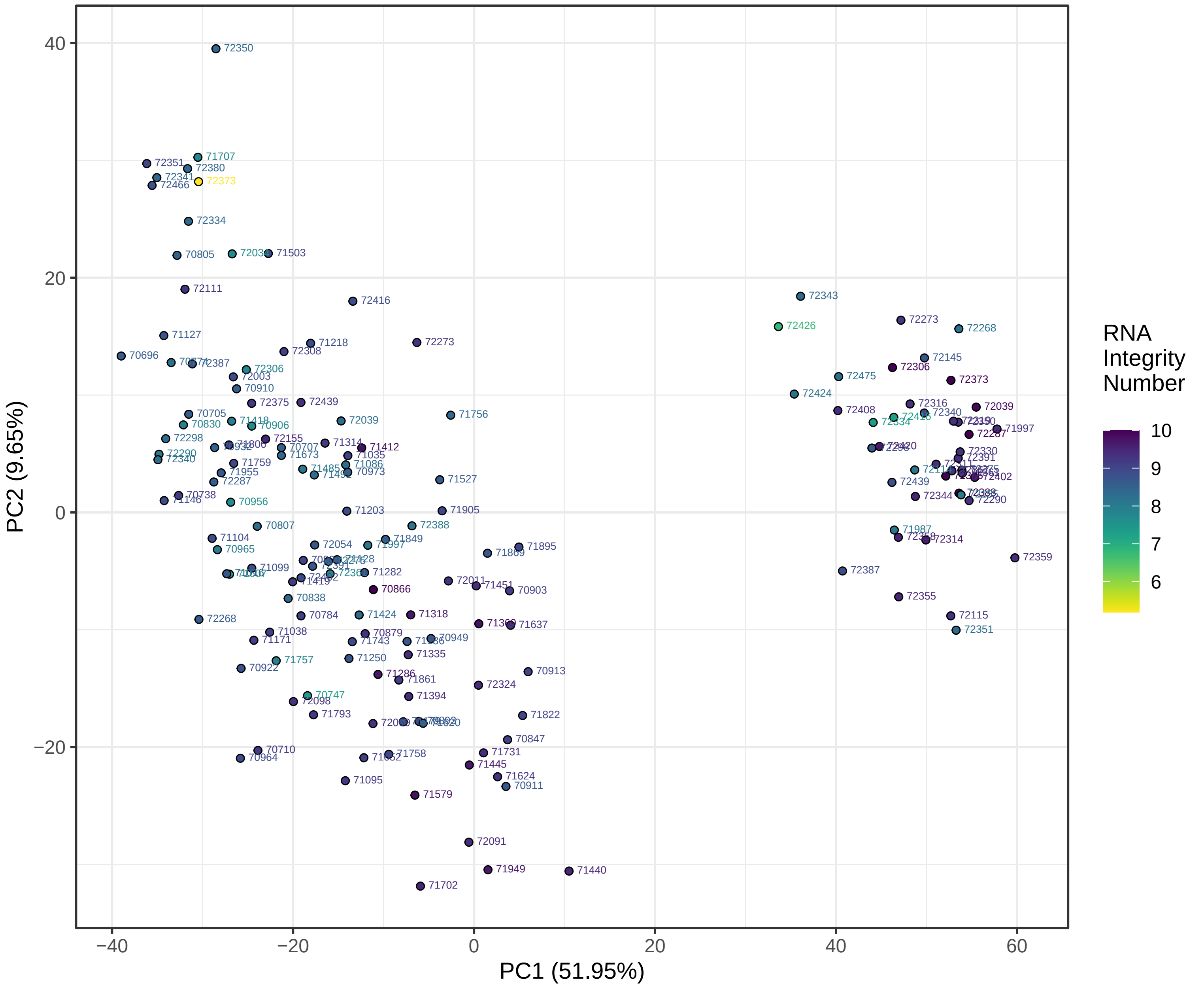

**Table S1: Demographics and Pregnancy Outcomes of All Sequenced Subjects, with the Matched Cohort Analyzed as a Third Group**

| | T1 Only | T3 Only | Matched Group | P value |
|---|---|---|---|---|
| **N** | 101 | 20 | 23 | |
| **Parental ages** | | | | |
| **Maternal age (SD)** | 37.8 (2.8) | 37.1 (3.4) | 36.5 (2.7) | 0.17 |
| **Paternal age (SD)** | 39.6 (4.7) | 39.8 (5.1) | 36.2 (3.1) | 0.0015<br>1st vs. match = 0.0008<br>3rd vs. match = 0.0016 |
| **Maternal race/ethnicity (%)** | | | | |
| Caucasian | 99 (98.0%) | 14 (70%) | 20 (87.0%) | <0.001<br>1st vs. match = 0.04<br>3rd vs. match = 0.26 |
| Caucasian<br>Asian<br>African American<br>Biracial (Cauc/Asian)<br>Biracial (Other) | 99 (98%)<br>1 (0.99%)<br>1 (0.99%)<br>0 (0%)<br>0 (0%) | 14 (70%)<br>2 (10%)<br>2 (10%)<br>1 (5%)<br>1 (5%) | 20 (87.0%)<br>1 (4.4%)<br>0 (0%)<br>1 (4.4%)<br>1 (4.4%) | 0.001 |
| Non-Hispanic | 98 (97.0%) | 19 (95.0%) | 16 (69.6%) | <0.001<br>1st vs. match = <0.001<br>3rd vs. match = 0.050 |
| **Paternal race/ethnicity (%)** | | | | |
| Caucasian | 97 (96.0%) | 15 (75%) | 20 (87.0%) | 0.005<br>1st vs. match = 0.12<br>3rd vs. match = 0.44 |
| Caucasian<br>Asian<br>African American<br>Biracial (Cauc/Asian)<br>Biracial (Other) | 97 (96.0%)<br>1 (0.99%)<br>2 (1.98%)<br>1 (0.99%)<br>0 (0%) | 15 (75%)<br>2 (10%)<br>2 (10%)<br>1 (5%)<br>0 (0%) | 20 (87.0%)<br>0 (0%)<br>1 (4.4%)<br>0 (0%)<br>2 (8.7%) | 0.005 |
| Non-Hispanic | 99 (98.0%) | 19 (95.0%) | 17 (73.9%) | 0.001<br>1st vs. match <0.001<br>3rd vs. match = 0.10 |
| **Fetal race/ethnicity (%)** | | | | |
| Caucasian | 97 (96.0%) | 12 (60%) | 18 (78.3%) | <0.001<br>1st vs. match = 0.011<br>3rd vs. match = 0.32 |
| Caucasian<br>Asian<br>African American<br>Biracial (Cauc/Asian)<br>Biracial (Other)<br>Multiracial | 97 (96.0%)<br>1 (0.99%)<br>1 (0.99%)<br>1 (0.99%)<br>1 (0.99%)<br>0 (0%) | 12 (60%)<br>0 (0%)<br>2 (10%)<br>5 (25%)<br>0 (0%)<br>1 (5%) | 18 (78.3%)<br>0 (0%)<br>0 (0%)<br>2 (8.7%)<br>1 (4.4%)<br>2 (8.7%) | <0.001 |
| Non-Hispanic | 98 (97.0%) | 19 (95%) | 14 (60.9%) | <0.001 |
| **Fetal sex: female** (%) | 50 (49.5%) | 9 (45%) | 9 (39.1%) | 0.65 |

| | T1 Only | T3 Only | Matched Group | P value |
|---|---|---|---|---|
| **N** | 101 | 20 | 23 | |
| **Maternal pre-pregnancy BMI, kg/m$^2$ (SD)** | 21.7 (3.6) | 24.0 (5.1) | 23.0 (3.6) | 0.03<br>1st vs. match = 0.07<br>3rd vs. match = 0.82 |
| **Maternal pre-existing medical conditions** | | | | |
| Hypertension | 0 | 0 | 0 | |
| Diabetes | 0 | 0 | 0 | |
| Thyroid disorder | 3 (2.97%) | 6 (30%) | 0 (0%) | <0.001<br>1st vs. match = 1<br>3rd vs. match = 0.006 |
| Other | 2 (1.98%) | 2 (8.7%) | 1 (5%) | 0.16 |
| **Hypertension management in pregnancy** | | | | |
| Anti-Hypertensives | 0 (0%) | 3 (15%) | 1 (4.4%) | 0.002<br>1st vs. match = 0.19<br>3rd vs. match = 0.32 |
| Any Magnesium use (Ante- or Postpartum) | 0 (0%0 | 5 (25%) | 2 (8.7%) | <0.001<br>1st vs. match = 0.03<br>3rd vs. match = 0.22 |
| **Pregnancy complications** | | | | |
| Hypertension (not pre-existing) | 0 (0%) | 6 (30%) | 3 (13.0%) | <0.001<br>1st vs. match = 0.006<br>3rd vs. match = 0.26 |
| Diabetes (not pre-existing) | 4 (3.96%) | 0 (0%) | 0 (0%) | 1.0 |
| Coagulation d/o | 0 | 0 | 0 | |
| Placenta previa | 0 (0%) | 0 (0%) | 1 (4.4%) | 0.30 |
| Placental abruption | 0 | 0 | 00 | |
| Placenta other | 0 (0%) | 1 (5%) | 1 (4.4%) | 0.09 |
| **Delivery** | | | | |
| Mode of delivery: Cesarean section, N (%) | 28 (27.7%) | 7 (35%) | 7 (30.4%) | 0.80 |
| Gestational age at delivery (SD), days | 276.2 (7.0) | 276.3 (8.7) | 276.8 (8.8) | 0.65 |
| Birthweight (SD), grams | 3420.8 (441.2) | 3599.6 (515.6) | 3393.0 (456.1) | 0.24 |

*Numbers are formatted as “mean (standard deviation)” or “count (percentage)”. N= count. SD = standard deviation. P-values are a three-way analysis, with additional two-way analysis shown for significant results. “1st”, “3rd” and “match” refer to the three groups.

**Figure S2A: Quality control heatmap, first trimester placenta.**

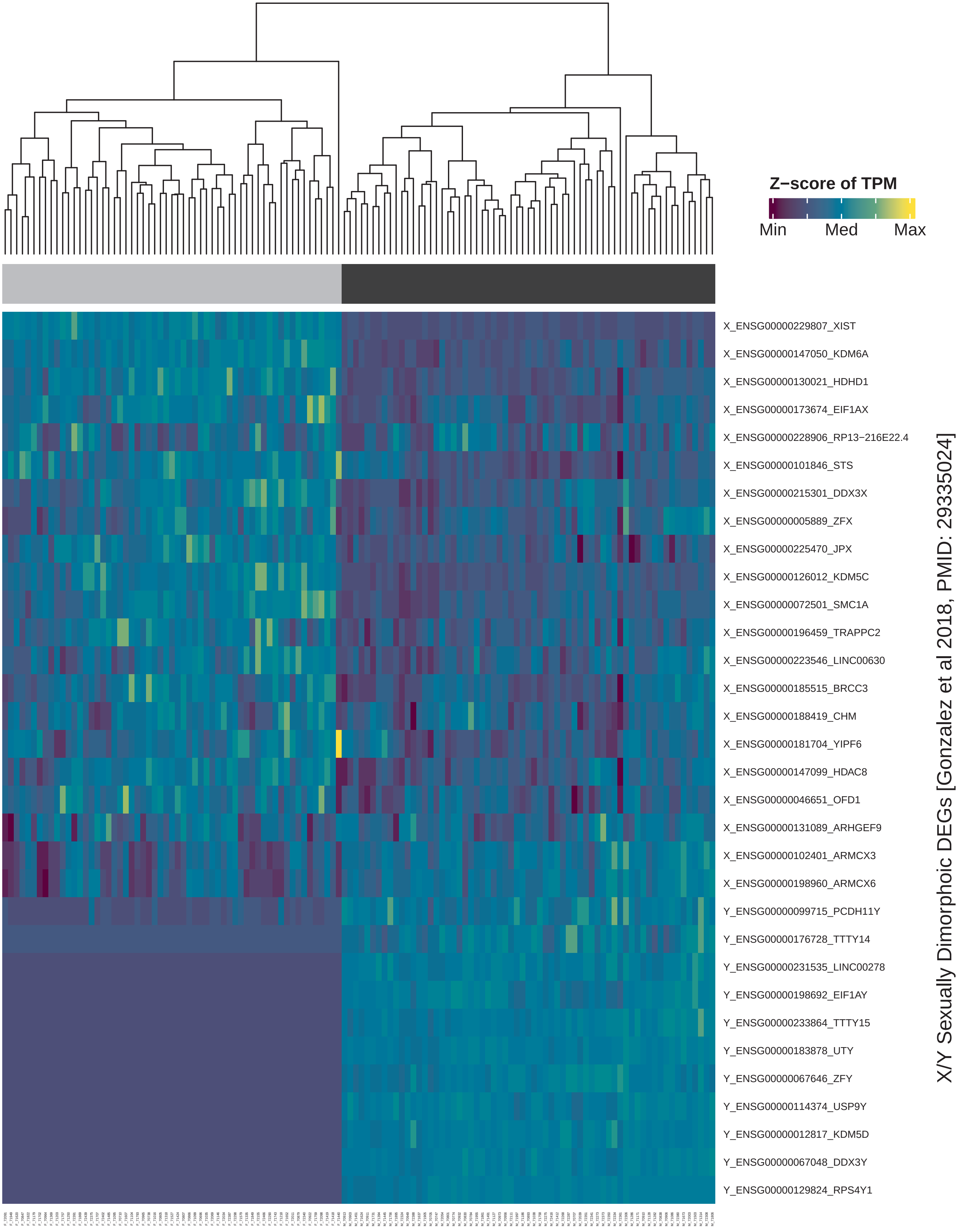


First trimester placenta samples sort as expected by fetal sex

**Figure S2B: Quality control heatmap, third trimester placenta.**

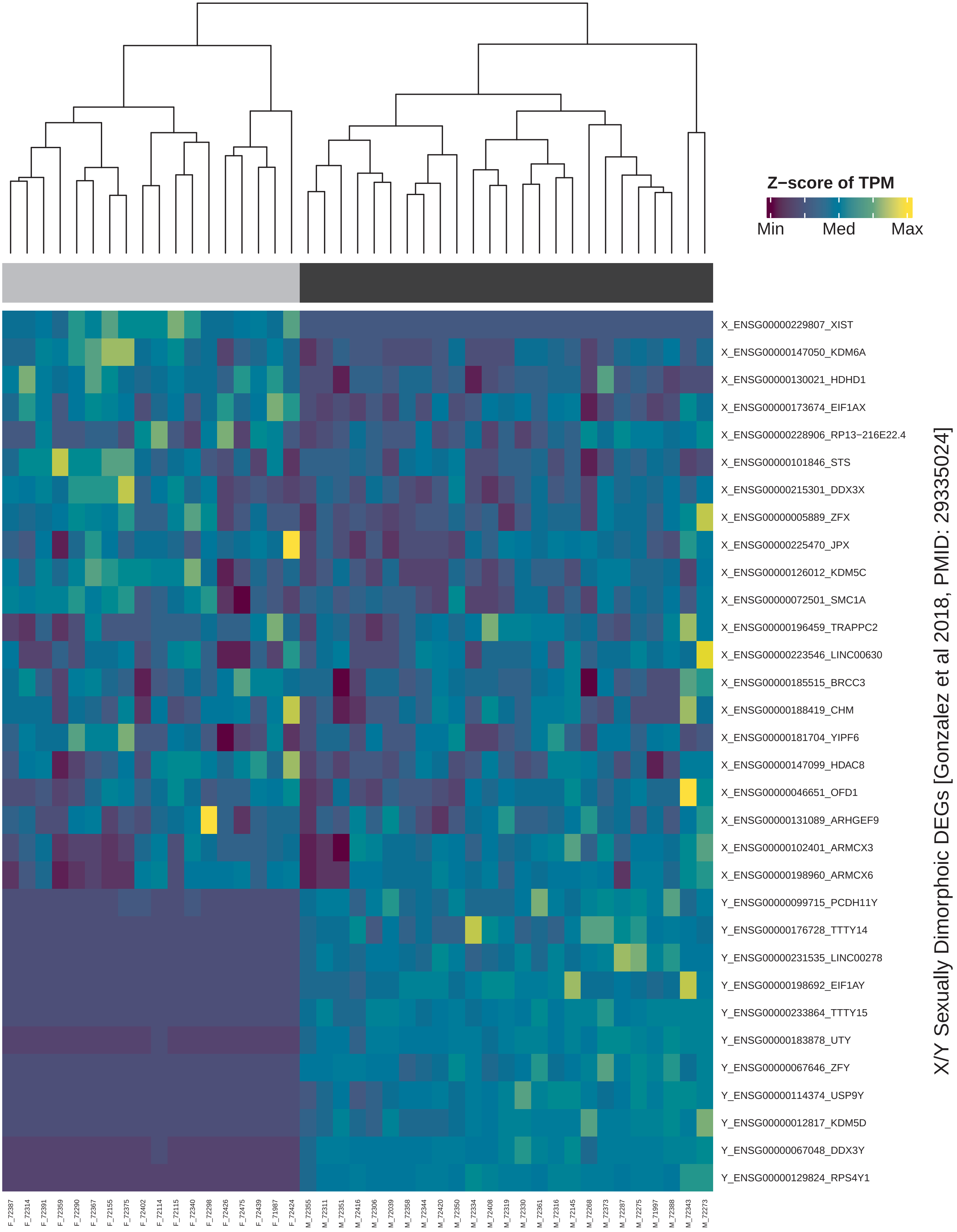


Third trimester placenta samples sort as expected by fetal sex

### Figure S3: Selection of the Study−Specific Expression Threshold, TPM>0.66

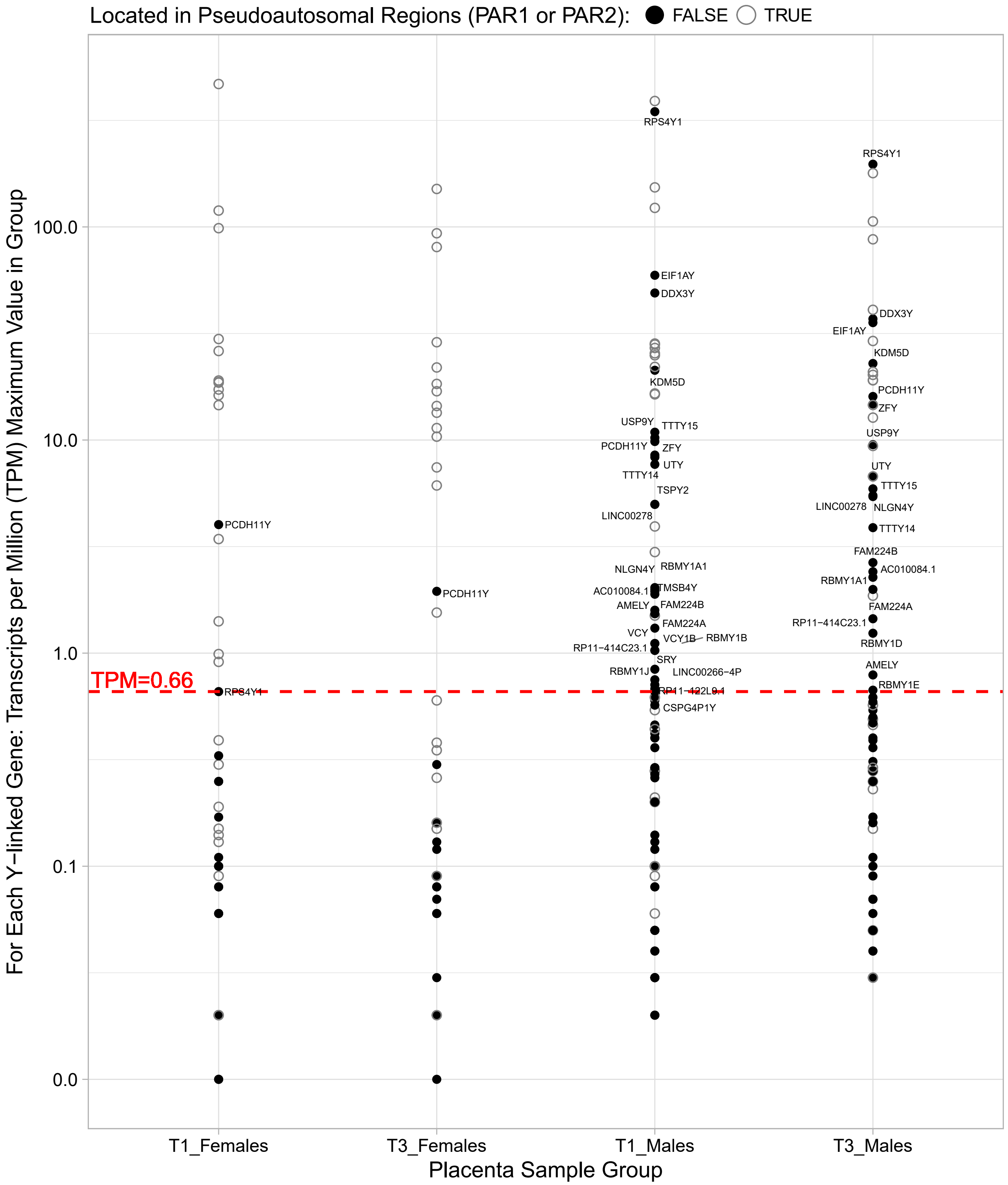


*Non−pseudoautosomal region genes with TPM>0.65 are labeled. T1 = first trimester. T3 = third trimester. PCDH11Y is not located in PAR1 or PAR2 but originates from a recent duplication from chromososome X and is more similar to its X−linked homolog than other X/Y homolog pairs. PCDH11Y signal in females is likely from PCDH11X.*

### Figure S4: Pathway enrichment analysis for the top 15% expressed protein coding genes in human placenta.

**A**

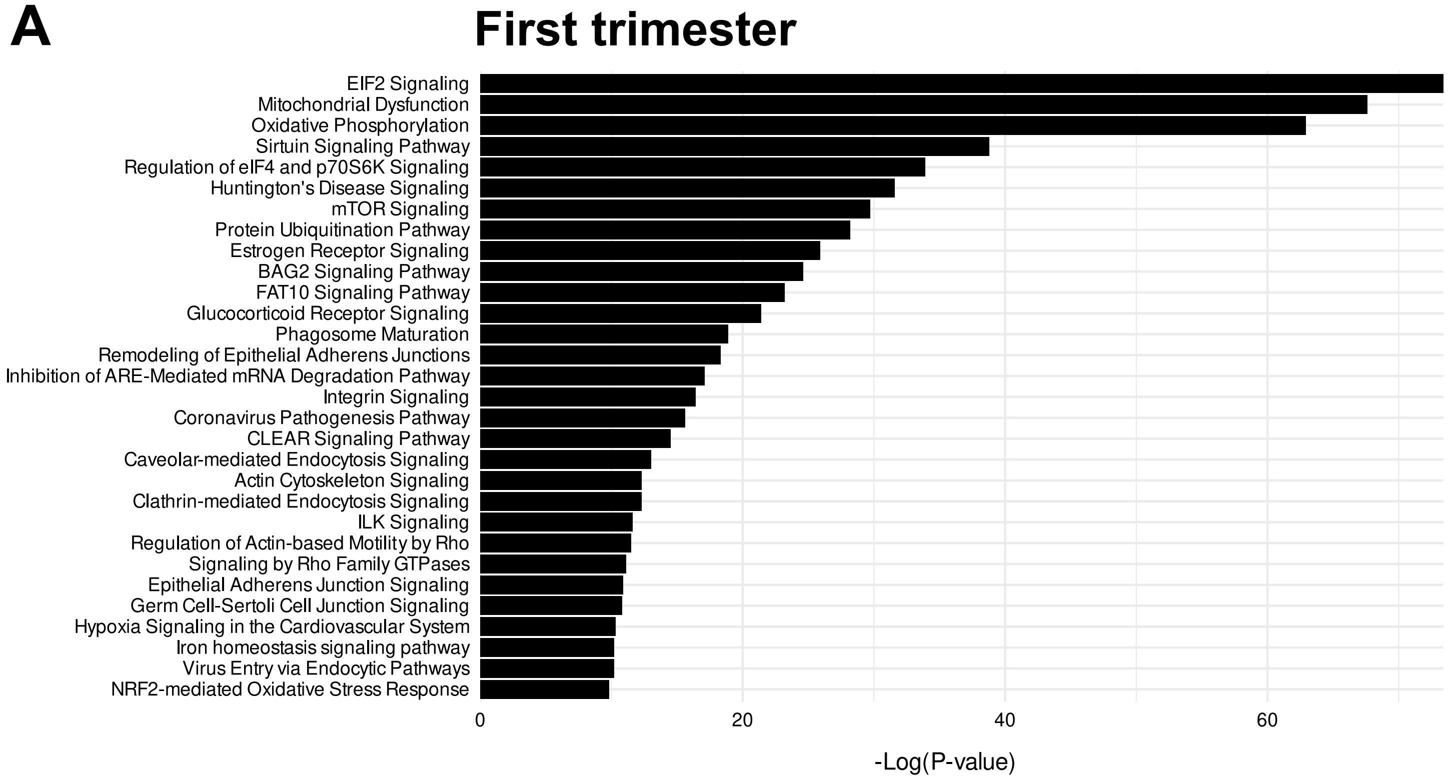


**B**

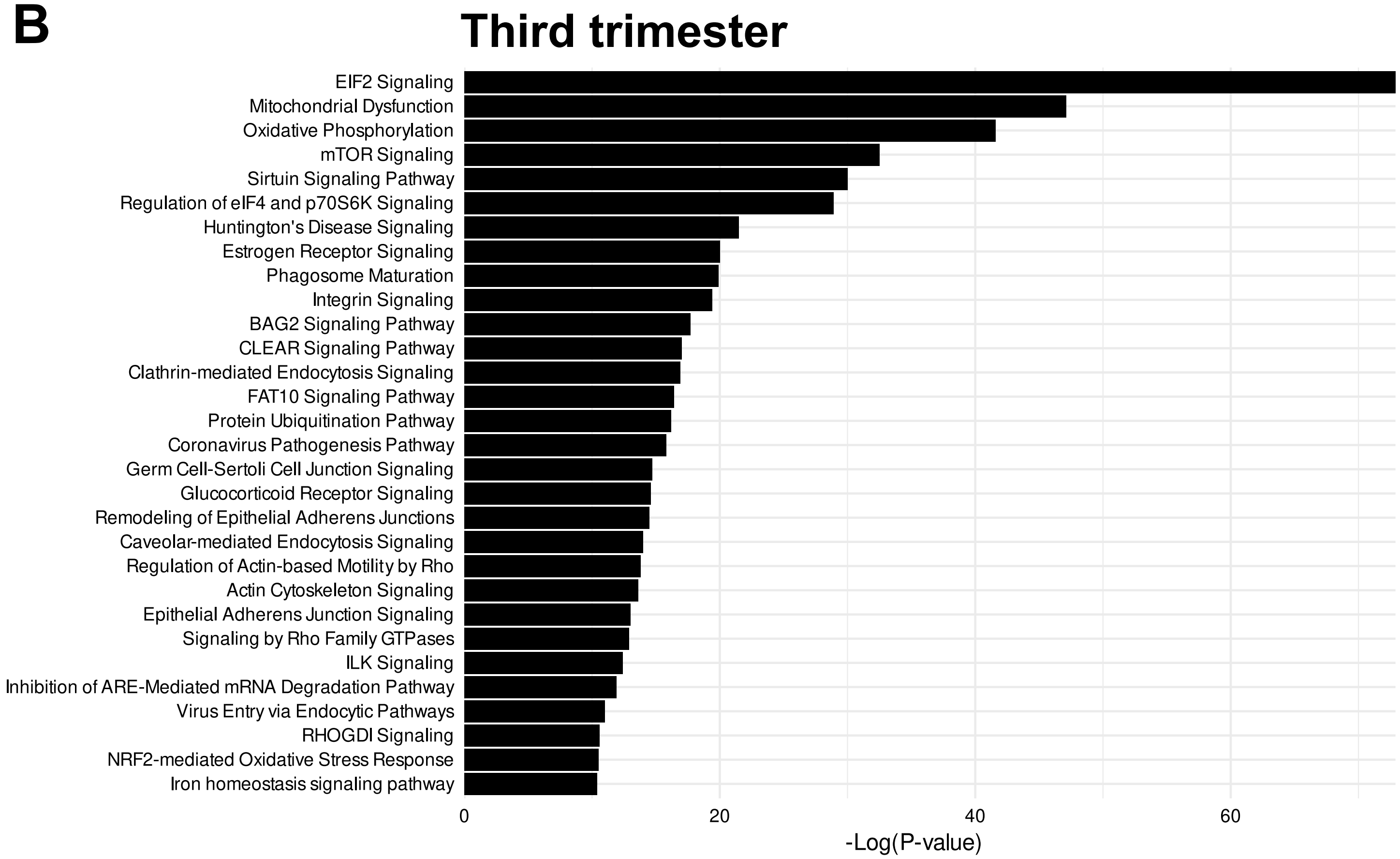

### Figure S5: Selection of strict DEG thresholds.

**A** All DEGs, either analysis:
TPM>0.66, FDR<0.05

**B** Strict DEGs, either analysis:
TPM>0.66, FDR<0.001, FC>1.5

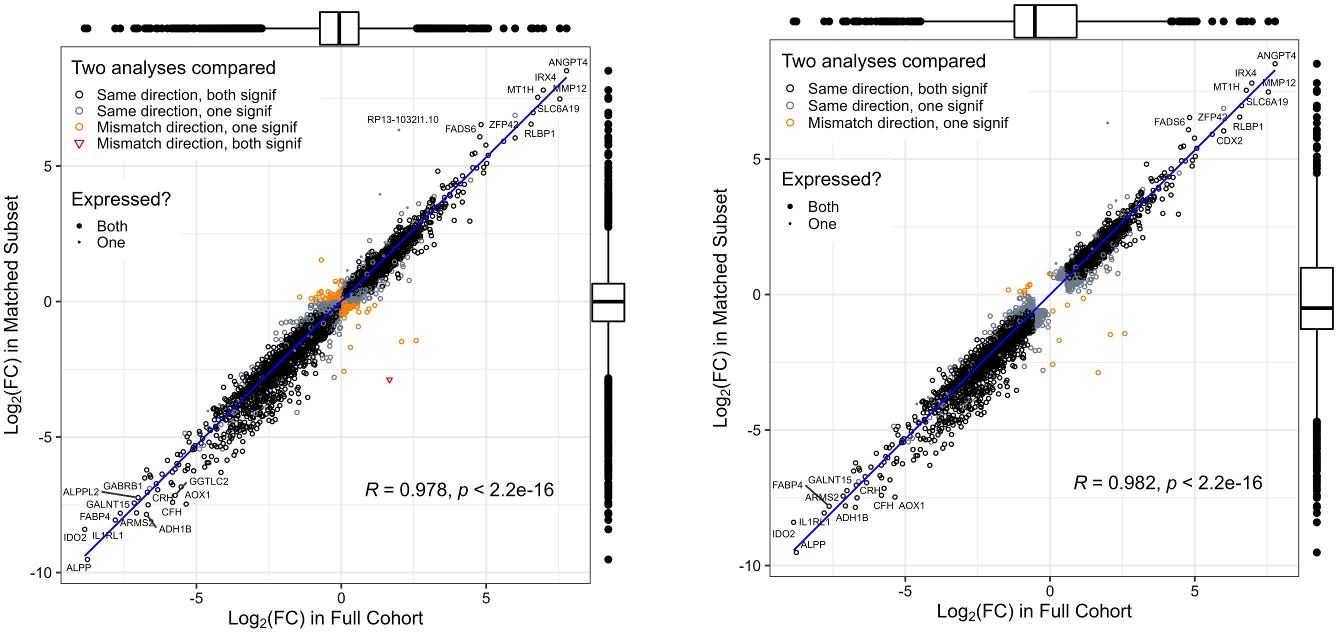


**A. All differentially expressed genes (DEGs) in either analysis.**
FDR<0.05, TPM>0.66 either trimester, protein coding
Total 12,850 protein coding genes

**B. All strict DEGs in either analysis.**
FDR<0.001, FC>1.5, TPM>0.66 either trimester, protein coding
Total 7,673 protein coding genes

Log2(FC) is the log-transformed fold change of first trimester vs third trimester
Positive Log2(FC) means higher expression in first trimester
Negative Log2(FC) means higher expression in third trimester

Box and whisker plots summarize the results for each axis (full or matched analysis)

*R* = Pearson correlation coefficient

*p* = linear regression p-value
